# SECE: accurate identification of spatial domain by incorporating global spatial proximity and local expression proximity

**DOI:** 10.1101/2023.12.26.573377

**Authors:** Yuanyuan Yu, Yao He, Zhi Xie

## Abstract

**Motivation:** Accurate identification of spatial domains is essential for analyzing spatial transcriptomics data to elucidate tissue microenvironments and biological functions. Existing methods utilize either local or global spatial relationships between spots to aid domain segmentation. A method that can concurrently capture both local and global spatial information may improve identification of spatial domains.

**Results:** In this article, we propose SECE, a deep learning-based method that captures both local and global relationships among spots and aggregates their information using expression similarity and spatial similarity. We benchmarked SECE against eight state-of-the-art methods on six real spatial transcriptomics datasets spanning four different platforms. SECE consistently outperformed other methods in spatial domain identification accuracy. Moreover, SECE produced spatial embeddings that exhibited clearer patterns in low-dimensional visualizations and facilitated more accurate trajectory inference.

**Availability and implementation:** SECE is implemented and provided as a pip installable Python package which is available on GitHub https://github.com/xie-lab/SECE.

## 1 Introduction

Spatial transcriptomics (ST) captures gene expression profiles with spatial information, providing novel insights into tissue molecular heterogeneity. Applying ST technology plays a crucial role in identifying cell-cell interactions, signaling pathways within the tissue microenvironment, and has enabled groundbreaking discoveries across fields such as neuroscience (Maynard, et al., 2021), developmental biology (Wang, et al., 2022), and cancer biology (Hunter, et al., 2021). A variety of ST platforms have been developed with varying throughput and resolution. Image-based ST platforms, including STARmap (Wang, et al., 2018), seqFISH (Lubeck, et al., 2014; Shah, et al., 2016), seqFISH+ (Eng, et al., 2019), MERFISH (Moffitt, et al., 2018) and FISSEQ (Lee, et al., 2014), provide highly accurate gene expression measurement at single cell resolution but only for a limited number of targeted genes (Moses and Pachter, 2022). On the other hand, sequencing-based ST platforms, such as Spatial transcriptomics (Stahl, et al., 2016) and its commercial version 10X Genomics Visium, Slide-seqV2 (Stickels, et al., 2021), HDST (Vickovic, et al., 2019), Seq-Scope (Cho, et al., 2021) and Stereo-seq (Chen, et al., 2022), can perform high-throughput sequencing on a genome-wide scale with increasing spatial resolution. The spatial resolution of sequencing-based technology continues to improve, with Seq-Scope and Stereo-seq capable of merging subcellular spots based on cellular location to achieve single-cell resolution.

In ST data, spatial domains refer to regions exhibiting consistent patterns in both gene expression and physical location, each with specific anatomical structures (Maynard, et al., 2021; Ortiz, et al., 2020). Accurately identifying spatial domains is crucial for various downstream analyses, including trajectory inference, cell type deconvolution, and cell-cell communications, as well as their biological interpretation. Spatial domains are distinct from cell types that have been extensively studied in single-cell data. Cell types can be obtained by clustering transcriptional information, and their spatial distribution patterns are uncertain. They may be spatially concentrated, such as excitatory neurons, or discretely distributed, such as astrocytes (Zeisel, et al., 2018). Spatial domains are continuous in space, so relying solely on gene expression is insufficient to capture them. Integrating spatial location information is necessary to ensure continuity of domains, which presents a new challenge.

Several methods have been developed to address this challenge. Many existing techniques utilize spatial location information to find neighboring spots for each spot and enhance the similarity between neighbors to ensure the spatial continuity of the domain. Among them, BayesSpace (Zhao, et al., 2021) and BASS (Li and Zhou, 2022) perform latent variable modeling of regional labels and use Bayesian methods for inference. SpaGCN (Hu, et al., 2021) and STAGATE (Dong and Zhang, 2022) employ graph convolutional networks and graph attention networks to aggregate neighbor information, respectively. GraphST (Long, et al., 2023), SpaceFlow (Ren, et al., 2022), and conST (Zong, et al., 2022) utilize self-supervised graph embedding learning strategies. However, spatial adjacency relationships only represent local information, neglecting to consider the global structure and patterns may result in a lack of comprehensive understanding of the data. In contrast, SpatialPCA (Shang and Zhou, 2022) introduces a spatially aware dimension reduction method, which leverages global spatial relationship by measuring pairwise spots similarity. Nonetheless, focusing solely on global similarities may overlook subtle spatial details. Moreover, the simultaneous capture of both local and global information, harnessing the advantages of each, remains an area needing further exploration.

To this end, we developed SECE, an accurate method for identifying spatial domains in ST data by incorporating both global and local proximity. Global proximity is quantified by physical distance, and it is considered that the pairwise similarity between spots decreases with longer spatial distances. Local proximity, on the other hand, is determined by expression similarity, aggregating neighbor information based on the similarity of gene expression between spots. SECE utilize graph neural networks for spatial embedding (SE) learning, it aggregates local expression similarity through an attention mechanism while simultaneously constraining the global spatial similarity using a Gaussian kernel function. Subsequently, by performing clustering on the SE, we can derive the spatial domain to which each spot belongs. SE also facilitate downstream analyses like visualization and trajectory inference. We demonstrated SECE’s versatility across diverse ST platforms, including high-resolution methods like STARmap/Slide-seqV2/Stereo-seq and lower-resolution platforms like Visium. SECE’s accurate spatial representations in brain and tumor datasets highlight its ability to gain biological insights from complex ST data.

## 2 Materials and Methods

SECE is versatile for modeling ST data across resolutions including subcellular, single-cell, near-single-cell, and multicellular (Fig. 1A). It takes gene expression matrix and spatial coordinate matrix of ST data as input, and outputs spatial domains and embeddings for each spot. First, SECE uses an autoencoder (AE) module with count distribution assumption to compress the expression matrix into expression features. Next, it converts spatial coordinates into local and global position relationships, storing them in the adjacency matrix (ADM) and spatial similarity matrix (SSM), respectively. Then, the graph attention network (GAT) module is utilized to balance the expression similarity of local neighbors and the global spatial similarity to obtain SE (Fig. 1B). Finally, spatial domains are identified by clustering SE using mclust (Scrucca, et al., 2016). Additionally, downstream analyses, including low-dimensional visualizations (McInnes, et al., 2018; Van der Maaten and Hinton, 2008) and trajectory inference (Cao, et al., 2019; Wolf, et al., 2019), are derived from SE (Fig. 1C).

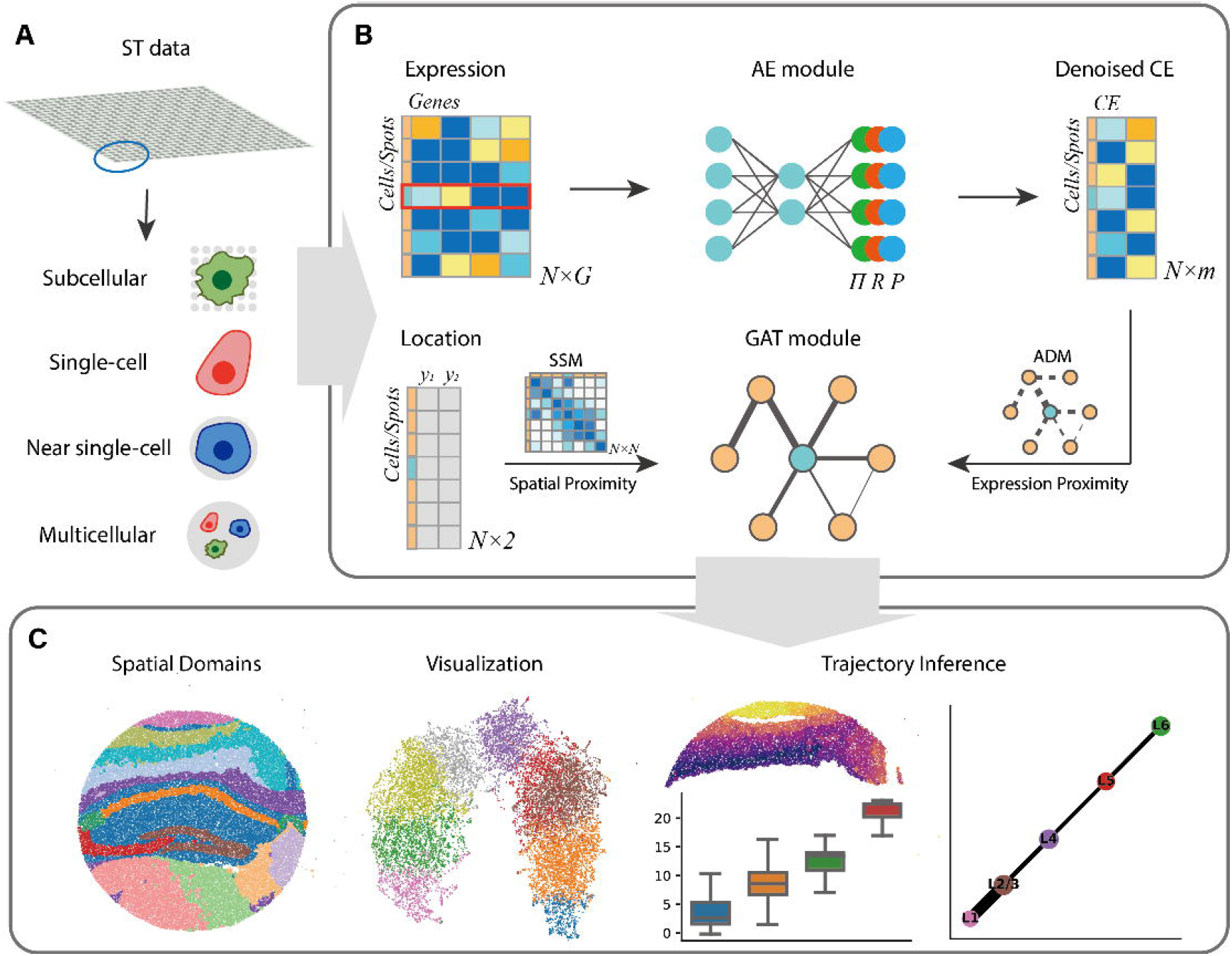

### 2.1 Extracting expression features with AE module

Given ST data with *N* spots and *G* genes, the dimensions of gene expression matrix *X* and spatial coordinate matrix *Y* are *N* x *G* and *N* x *2*, respectively. The raw counts *X*, are normalized by library size and then log-transformed to obtain the normalized expression matrix 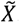. We first employ AE with zero-inflated negative binomial (ZINB) or negative binomial (NB) distribution (Eraslan, et al., 2019; Lopez, et al., 2018) to compress 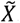 into low-dimensional features *Z*. Let *x_ng_* denotes the count value of gene *g* in spot *n*, the likelihood function of *x_ng_* under ZINB and NB distributions is given by:

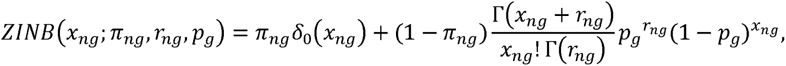

and

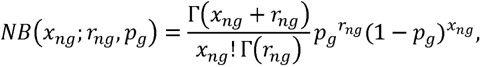

where *δ*_0_ is the Dirac delta function, *π_ng_* is the probability of true gene expression being 0, Γis the gamma function. (*r_ng_, p_g_*) is the standard parametrization for the NB distribution.

The AE module takes 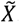, as input, and outputs distribution parameters. The formulation is:

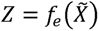

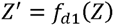

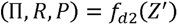

where *f_e_* represents an encoder, while *f_d1_* and *f_d2_* constitute the decoder. Specifically, *f_e_* consists of two nonlinear layers, each utilizing Rectified Linear Unit (*ReLU*) activation functions. These layers reduce the feature dimension *G* into m′ and m, respectively, yielding the expression feature *Z*. Subsequently, *f_d1_* decodes *Z* into *Z′* with a feature dimension of m′. *f_d2_* comprises three output layers, which take *Z′* as input and output three parameter matrices (Π,*R, P*) of the ZINB distribution, each consisting of elements (*π_ng_, r_ng_,p_g_*).The activation functions of these three output layers are exponential, sigmoid, and exponential functions respectively. *f_d2_* employs two layers to learn(*R,P*) under the NB distribution assumption.

The goal is to minimize the reconstructed loss by minimizing the negative log-likelihood (NLL) function, that is, 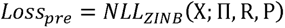 or 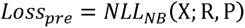. We employed ZINB for highly sparse data like Stereo-seq and Slide-seq, and NB for less sparse data including STARmap and Visium. Both are implemented in SECE package.

### 2.2 Capturing both local and global relationships

To incorporate physical location information, we construct an ADM and SSM from the spatial coordinates *Y*. ADM summarizes local spatial relationships by storing neighbors for each spot. This local neighborhood information is later employed to adaptively aggregate features in GAT module based on expression similarity. In contrast, SSM captures global spatial proximity by providing a spatial similarity measure for all pairs of spots, not just neighbors. The SSM is subsequently utilized to constrain the global similarity of SE.

ADM *A* is a *N*×*N*-dimensional symmetric matrix, where elements are assigned values of 1 or 0 to indicate neighboring spots or non-neighboring spots, respectively. More precisely, the element *A_ij_* denotes the adjacent relationship between spot *i* and spot *j*:

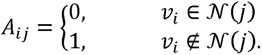

Here *N*(*j*) represents the set of spatial neighbors of spot *j*, which can be determined based on coordinates *Y* by employing K-Nearest Neighbor (KNN) or applying a distance cutoff. By default, we utilize KNN, with the number of neighbors set to 6 for Visium datasets and 10 for other datasets.

SSM Σ is also a *NXN*-dimensional symmetric matrix, wherein elements decrease as the distance between spots increases, exhibiting an exponential decay tendency. For spot *i* and spot *j* with coordinates *y_i_* and *y_j_*, the correspond element is expressed as:

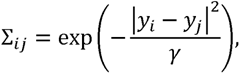

where the bandwidth parameter *γ* controls the spatial influence. By default, *γ* is set as the 0.05 quantile distance. A larger *γ* results in a greater spatial influence.

### 2.3 Learning SE with GAT module

After capturing the expression features and extracting local and global position relationships, we employ the GAT module to learn SE, subsequently clustering SE to delineate spatial domains. The GAT module consists of two GAT layers.

We first introduce the GAT layer. It takes feature matrix and ADM as input, and outputs a new feature matrix after aggregating neighbor information. Let 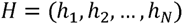 denotes the input feature matrix, which has dimensions 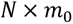, with *N* samples and *m*_0_ features. The output of GAT layer is denoted as 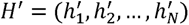 with dimensions 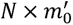. The GAT layer performs aggregation for each sample adaptively based on the normalized attention scores.

For sample *j* the output feature 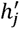 can be formulated as layer performs aggregation for each sample adaptively based on the normalized attention scores.

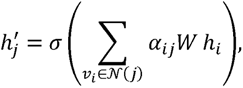

where *W* is a weight matrix with dimensions 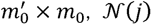 represents the set of neighboring samples of sample *j*, *α_ij_* is the normalized attention coefficient matrix using the SoftMax function:

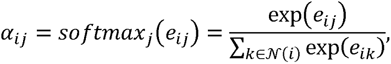

where 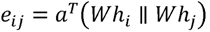, *α* is learnable vector and 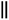 is the concatenation operation. We used the Exponential Linear Unit (ELU) as activation function *σ* in the GAT layer.

The GAT module in SECE consists of two GAT layers. It takes expression features *Z* and ADM *A* as input, and outputs SE matrix *U*, which is a *NXm*-dimensional matrix:

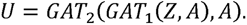

During neighbor information aggregation using GAT, local expression similarities are captured via attention and adaptively aggregated. To preserve as much information as possible in expression features, the local learning target is the reconstruction loss 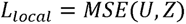. We further constrain pairwise correlation using SSM that including global information, that is, the correlation of each SE at, positions *UU^T^*, is close to Σ, 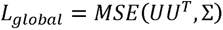. The objective function balances the two similarities by *λ_global_* and *λ*_local_:

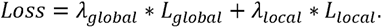

We kept *λ*_local=1_ unchanged and adjusted *λ_global_* based on the ST platform. After obtaining the final SE *U*, we utilized the clustering method mclust (Scrucca, et al., 2016) to cluster the *U* and determine the spatial domain for each spot.

### 2.4 Architecture of SECE

In the AE module, the dimension of the hidden layer *m*′ and bottleneck layer are 128 and 32, respectively. Adaptive Moment Estimation (Adam) optimizer is used to minimize *Loss_pre_* with a learning rate of 1e-3 and dropout rate of 0.1. For the GAT module, the dimensions of the two GAT layers are both 32. The Adam optimizer is employed to minimize *Loss*with a learning rate of 1e-2 and dropout rate of 0.2. The default number of iterations for the AE and GAT modules are set to 40 and 50, respectively.

The hyperparameters *λ_global_* and *λ*_local_ control the contributions of global and local similarities, respectively, and a larger *λ_global_* giving greater global influence. Each ST platform has different resolutions and fields of view. For example, spots of ST arrays contain dozens of cells, while Stereo-seq only contain single cell. Stereo-seq can sequence the hemibrain while STARmap can only detect a minor region of the visual cortex. We choose smaller *λ_global_* value for platforms with more spots and higher resolution. Specifically, for Stereo-seq and Slide-seqV2 data including over 10,000 spots with approximately single-cell resolution, we used *λ_global_* = 0.08. For Visium data with several thousand spots *λ_global_* was set to 0.3. For STARmap data with only 1207 cells, we set *λ_global_* to 2. Users could also select *λ_global_* following this standard.

## 3 Results

### 3.1 Application to STARmap data

We first tested SECE on the mouse visual cortex data generated by STARmap (Wang, et al., 2018) with ground truth layers. The 1,207 cells were divided into 7 layers, including Layer(L)1, L2/3, L4, L5, L6, as well as the corpus callosum (CC) and hippocampus (HPC) (Fig. 2A).

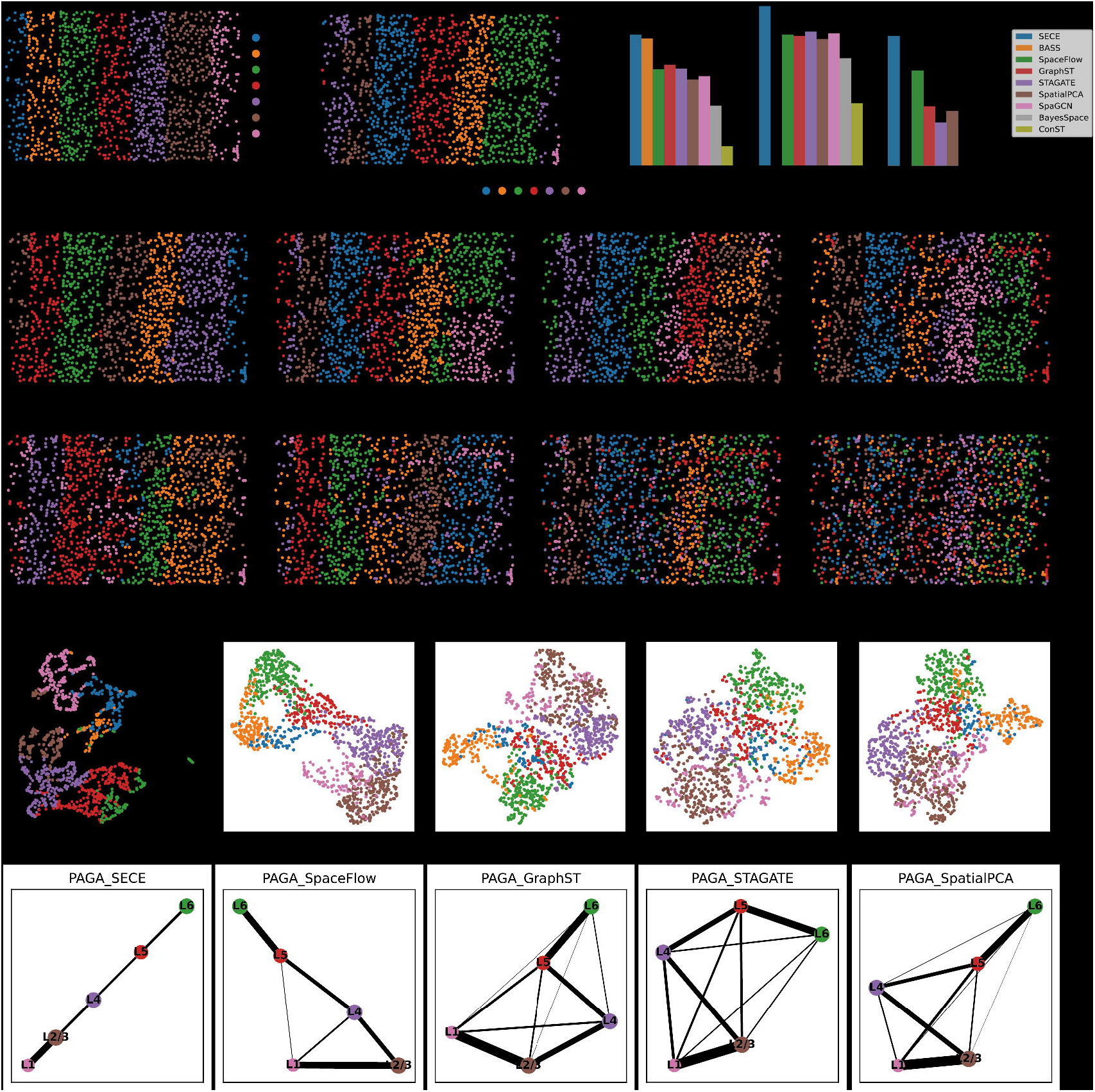

First, we compared the spatial domain identification results of SECE with eight existing methods, including BASS, SpaceFlow, GraphST, STAGATE, SpatialPCA, SpaGCN, BayesSpace, and conST (Fig. 2B, C). The consistency of spatial domains with the ground truth layers were evaluated using Adjusted Rand Index (ARI) and Accuracy (ACC) based on Kuhn-Munkres (Munkres, 1957). SECE achieved the highest consistency, with an ARI value of 0.65 and an ACC value of 0.79 (Fig. 2D). It was closely followed by BASS with ARI of 0.63, however, BASS incorrectly combined HPC and L5 (domain 5), which prevents the utilization of the Kuhn-Munkres algorithm to reassign clusters and compute the accuracy (ACC). Furthermore, other algorithms also exhibited weak spatial aggregation, as indicated by the Local inverse Simpson’s index (LISI) (Korsunsky, et al., 2019) (Supplementary Figure S1A). Additionally, these methods mistakenly classified aggregated endothelial cells (Endo) in L1 and L2/3 into the same domain, such as domain 2 in SpaceFlow, domain 3 in STAGATE and BayesSpace, domain 5 in GraphST and SpatialPCA, and domain in SpaGCN (Fig. 2C, Supplementary Figure S1B).

Next, we compared SE of SECE with SpaceFlow, GraphST, STAGATE, and SpatialPCA. Due to the difficulty of conST in generating clear spatial domains, its SE learning comparison was excluded. SE of SECE explained the ground truth layers most effectively (ASW=0.16), followed by SpaceFlow (ASW=0.12), GraphST (ASW=0.07), STAGATE (ASW=0.07), and SpatialPCA (ASW=0.05) (Fig. 2D, right). Moreover, SECE had a clearer and continuous pattern in UMAP visualization compared to the other methods (Fig. 2E). When comparing trajectory inference results, we selected the cortex part in ground truth, namely L1, L2/3, L4, L5 and L6. The PAGA(Wolf, et al., 2019) analysis based on SECE revealed a linear and continuous relationship between these layers (Fig. 2F). Conversely, SpaceFlow mistakenly made the connection between L1 versus L4 and L5, while the patterns in the other three methods were more unclear. For individual cells, SECE exhibited a sequential increase of pseudo-time from L6 to L1 (Supplementary Figure S1C, D) based on Monocle3(Cao, et al., 2019), and the similar trend was also observed in SpaceFlow. However, GraphST, STAGATE, and SpatialPCA ordered L1 and L2/3 incorrectly (Supplementary Figure S1E, F).

### 3.2 Application to Slide-seqV2 data

We further tested SECE on the hippocampal dataset generated by Slide-seqV2 platform (Stickels, et al., 2021). SECE identified 14 distinct domains, and we annotated them according to the known structure of Allen Brain Atlas (ABA) (Fig. 3A). The domains were hippocampus (Cornu Ammonis (CA)1, CA2, CA3, Dentate gyrus (DG), and CA slm/so/sr), cortex (Layers 4, 5a, 5b, 6), third ventricle, CC, and three subregions of the thalamus (Fig. 3B, left). We further meticulously verified the subtle spatial domains. For example, the four important components of the hippocampal region, CA1, CA2, CA3, and DG, were clearly demarcated, as evidenced by the high expression of their known markers *Wsf1*, *Rgs14*, *Nptxr* and *C1ql2* (Saunders, et al., 2018), respectively (Fig. 3C, Supplementary Figure S2A). Layer 5 in cortical regions was identified as two sublayers, Layer 5a and 5b, with different gene expression levels (Supplementary Figure S2A, B). The composition of cell types in each domain further supported the delineation (Supplementary Figure S3).

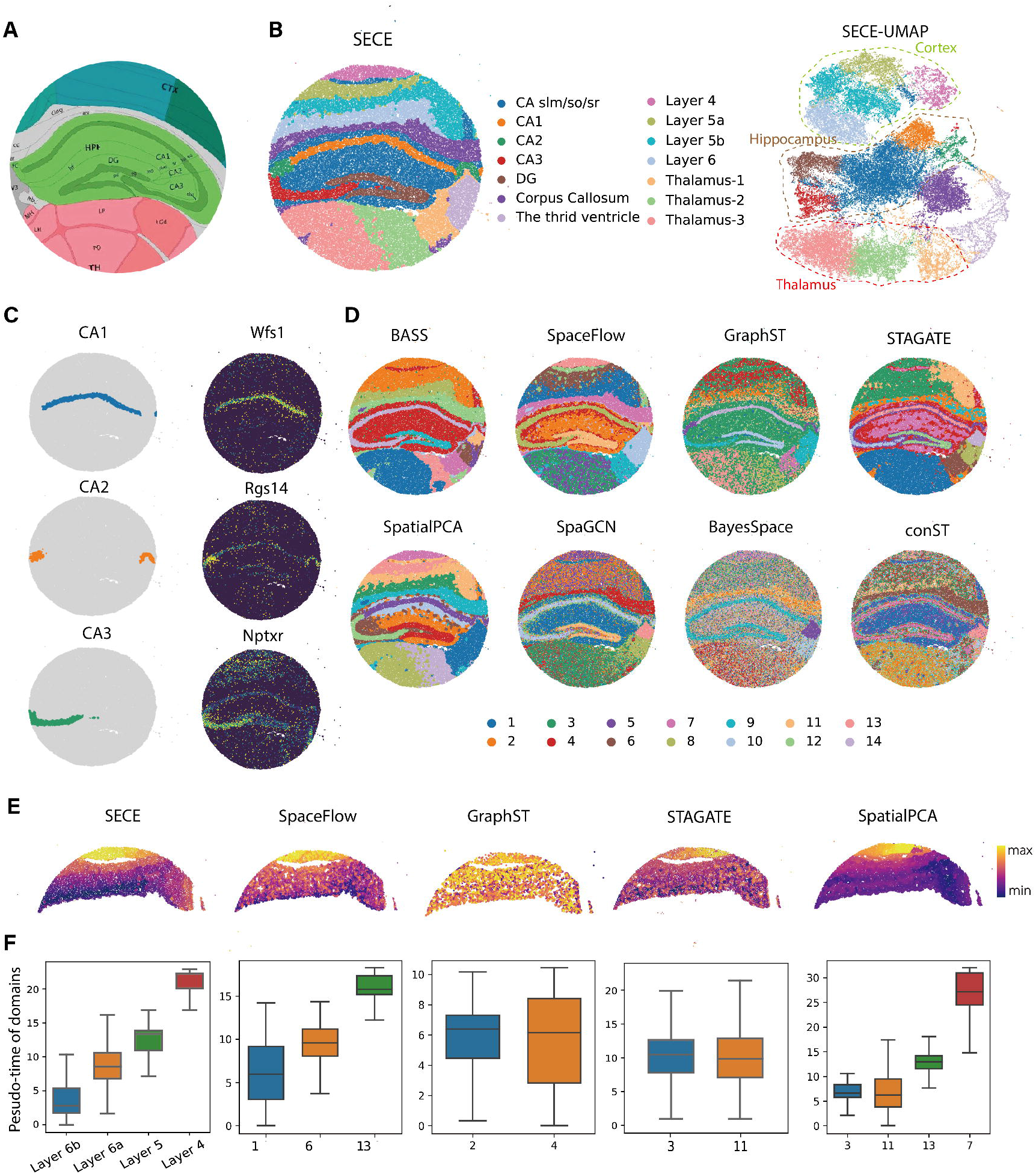

For comparison, we evaluated the performance of existing methods for spatial domain identification (Fig. 3D). Notably, SECE was the only approach that could accurately detect subregions in both the cortex and CA. Specifically, for cortical areas, BASS failed to distinguish between Layer 4 and Layer 5, SpaceFlow mixed Layer 5 and Layer 6, SpatialPCA exhibited suboptimal division smoothness, and the remaining algorithms were unable to generate clear cortical regions. In the case of CA, except for GraphST, none of the methods succeeded in identifying the CA2 region. Furthermore, SECE exhibited the highest spatial aggregation performance as it had the smallest LISI values (Supplementary Figure S4A).

We also compared the performance of SE. UMAP generated from SECE clearly displayed clustering patterns for the hippocampus, cortex, thalamus, third ventricle, and their subregions (Fig. 3B, right). In addition, the sublayers of them like Layer 4 and Layer 6 were arranged in a sequence, while UMAP of STAGATE and GraphST missed them (Supplementary Figure S4B). Moreover, we conducted trajectory inference for the cortex due to its continuous relationship between sublayers. We selected the domains corresponding to the cortex in each method and started with the deepest clusters, such as Layer 6 for SECE, domain 1 for SpaceFlow, domain 2 for GraphST, and domain 3 for SpatialPCA and STAGATE (Fig. 3E, F). For SECE, the pseudo-time consistently increased with decreasing cortical depth, clearly capturing the pseudo-time relationship between different layers. In contrast, SpaceFlow exhibited less distinguishable differences between the learned layers. SpatialPCA reversed the relationship between Layer 5b and Layer 6, while GraphST and STAGATE failed to identify the relationship within the cortical region.

### 3.3 Application to Stereo-seq data

In this section, we assessed on the mouse olfactory bulb (Fu, et al., 2021) Stereo-seq(Chen, et al., 2022) data. Spatial domains identified by SECE were annotated based on known olfactory bulb layers, including rostral migratory stream (RMS), granule cell layer (GCL), inner plexiform layer (IPL), mitral cell layer (MCL), external plexiform layer (EPL), glomerular layer (GL) and olfactory nerve layer (ONL) from inside to outside (Fig. 4A; Supplementary Figure S5A). These domains were validated using known markers (Saunders, et al., 2018) for each layer (Fig. 4B). Notably, besides the known seven layers, we made a finer division of GCL and ONL, and the sublayers were named GCL-Inner, GCL-Outer, ONL-Inner and ONL-Outer, respectively. Several evidences confirmed these sublayers. First, marker gene expression differed between them, where the GCL-inner highly expressed *Nrgn* and the GCL-outer highly expressed *Pcp4*. Markers in the ONL-outer also exhibited higher expression levels compared to those in the ONL-inner. Second, sublayers exhibited distinct cell type composition (Supplementary Figure S5B to D). GCL-Outer was almost composed of GC, while GCL-Inner contained a certain amount of Oligo, and the GC subtypes in GCL-Inner and GCL-Outer also differed. Moreover, the ONL-Outer contained almost exclusively OEC, while the ONL-Inner contained a fraction of OEC and more Astro. This supported previous findings that the ONL, as a part of the olfactory bulb blood-brain barrier, had fine internal and external subregions with different cell types (Beiersdorfer, et al., 2020). Our study provided further support for this finding at the spatially resolved single-cell level.

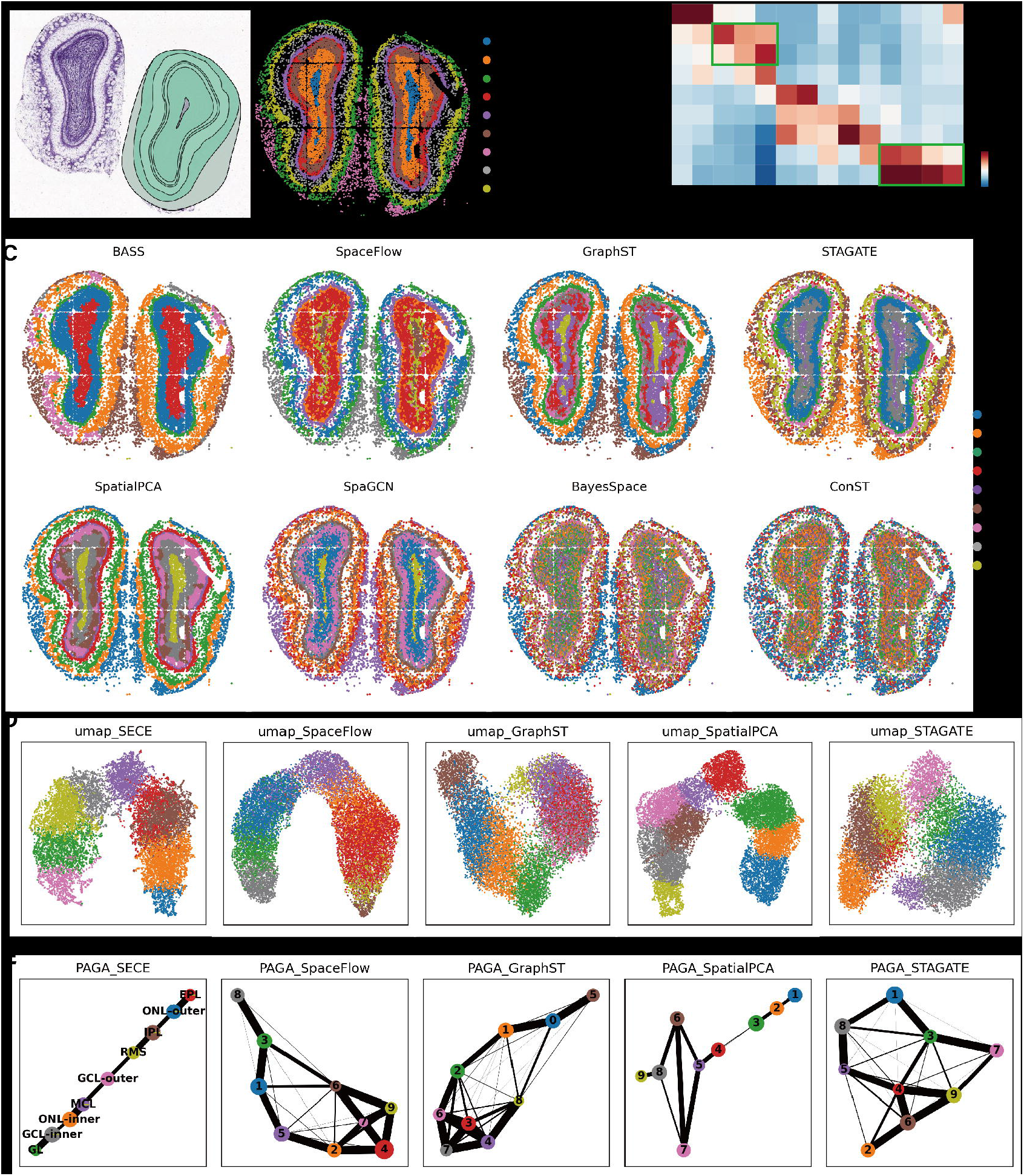

We also compared the performance of SECE to the existing methods (Fig. 4C). BASS failed to separate the RMS from the GCL layer and could not distinguish GL from EPL. It also identified some irrelevant domains with few cells, such as domain5, 8, and 9. STAGATE and SpaGCN mixed GL and EPL. SpaceFlow, GraphST, and SpatialPCA encountered challenges in accurately dividing the GCL layer. BayesSpace and conST did not yield a clear domain identification. Additionally, BASS had the most spatial aggregation performance with lowest LISI value, while SECE ranked second (Supplementary Figure S5E). Furthermore, SECE showed robustness when tested with different number of clusters (7, 8, and 10), consistently providing well-bounded ring stratification (Supplementary Figure S6). Moreover, we assessed the SE learning capabilities. The UMAP visualization based on SECE exhibited a continuous low-dimensional pattern with nine layers arranged in order of spatial position from inside to outside (Fig. 4D). The trajectory inference for these layers demonstrated approximately linear relationship (Fig. 4E). SpatialPCA also displayed linear patterns except for domains 6 and 7, while results of SpaceFlow, GraphST, and STAGATE exhibited many false positive connections between domains.

We further applied SECE onto mouse hemibrain dataset with a more complex anatomic structure (Chen, et al., 2022). SECE achieved spatial domain annotations that were in highly consistent with ABA anatomy (Fig. 5A, B, left). We could clearly separate different regions, including cortical regions, hippocampal regions, midbrain regions, thalamic regions, and fiber tracts (FT). The cortex included five layers: L1, L2/3, L4, L5, L6, LVC and CAA, which were supported by their cell type composition (Supplementary Figure S7, 8). In contrast, other methods failed to accurately identify these cortical layers (Fig. 5C, Supplementary Figure S9). Specifically, all of them encountered difficulties in identifying Cortex L5 with high smoothness. In addition, LISI showed that spatial domains identified by SECE had the strongest spatial pattern (Supplementary Figure S9A). Furthermore, low-dimensional visualization of SE using t-SNE (Van der Maaten and Hinton, 2008) showed the effect of aggregation in the same region and separation in different regions (Fig. 5B, right).

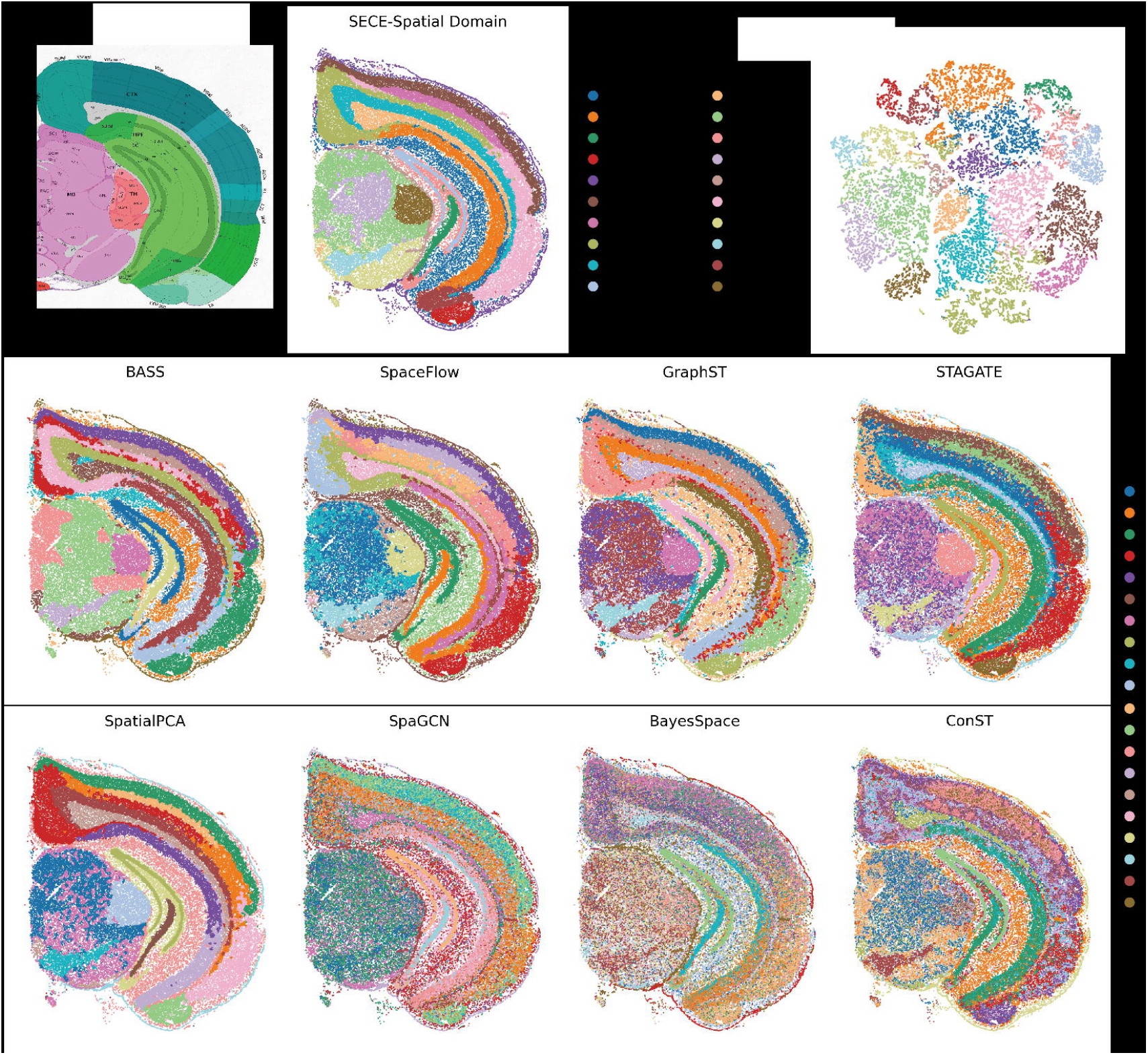

### 3.4 Application to Visium data

Finally, we tested the applicability of SECE on Visium dataset. The breast cancer dataset was divided into four phenotypic regions according to pathological images: ductal carcinoma in situ/lobular carcinoma in situ (DCIS/LCIS), healthy tissue (Healthy), invasive ductal carcinoma (IDC), and tumor surrounding regions with low features of malignancy (Tumor edge), as well as 20 annotated subdivisions (Fu, et al., 2021) (Fig. 6A; Supplementary Figure S10A).

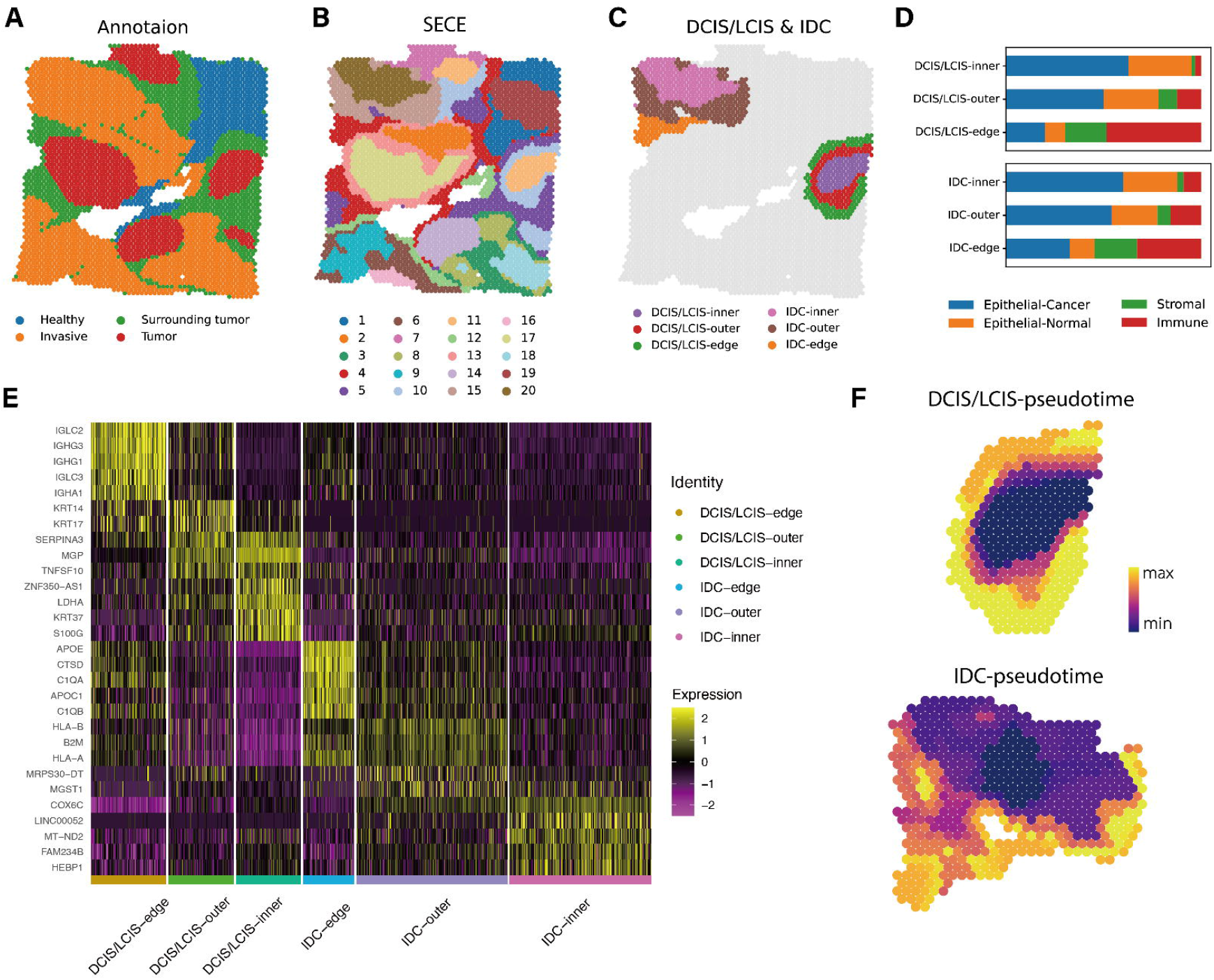

We initially segmented 20 domains using each algorithm and performed phenotype annotations, which were then compared with image-based manual annotations (Fig. 6B; Supplementary Figure S10B, C). We found that SECE divided individual tumor regions into wrapped layers more clearly. Specifically, cluster 20 and cluster 15 divided the IDC-5 into internal and external layers, while cluster 11 and cluster 10 split the DCIS/LCIS-1. (Fig. 6B; Supplementary Figure S10A). To investigate the biological significance of the refined stratification, we named clusters 20,15,11 and 10 as IDC-inner, IDC-outer, DCIS/LCIS-inner and DCIS/LCIS-outer, respectively. We also focused on the parts of clusters 4 and 5 that were located near the tumor edge, referred to as IDC-edge and DCIS/LCIS-edge. (Fig. 6C). We analyzed the cell type composition of each spot in these domains by integrating the annotated breast cancer scRNA-seq data (Wu, et al., 2021) and deconvoluting each spot using cell2location (Kleshchevnikov, et al., 2022) (Supplementary Figure S11, S12). The proportion of tumor cells gradually decreased as the tumor moves towards the edge, while the that of immune cells and stromal cells gradually increased (Fig. 6D), confirming the correctness of subregions.

We further explored the characteristics of these subregions (-edge,-outer and -inner) of DCIS/LCIS and IDC (Fig. 6E). Differential expression analysis(Hao, et al., 2021) showed that DCIS/LCIS-edge had a high level of humoral immunity, which could be confirmed by the enrichment of B cell receptor signaling pathway related genes (*IGLC2*, *IGLC3*, *IGHG1*, *IGHG3*, *IGHA1*). DCIS/LCIS-outer overexpressed *KRT14* of the keratin family, which was a key regulator of metastasis, suggesting invasive potential (Bilandzic, et al., 2019; Cheung, et al., 2016). In contrast, DCIS/LCIS-inner showed non-metastatic traits but also had tumor-promoting abilities, indicated by a high expression of *LDHA* (Martinez-Ordonez, et al., 2021). For the IDC subregions, IDC-edge showed elevated expression of biomarkers linked to tumor proliferation, invasion, and migration including tumor-associated macrophage (*APOE*), complement components (*C1qA*, *C1qB*), cathepsin (*CTSD*), and apolipoprotein (*APOC1*) (Nalio Ramos, et al., 2022; Revel, et al., 2022; Seo, et al., 2022; Zhang, et al., 2018; Zhang, et al., 2022). IDC-outer had a high level of immunity and some transferability, derived from the high expression of MHC class I-related genes (*HLA-A*, *HLA-B*, *B2M*) (Noblejas-Lopez, et al., 2019; Nomura, et al., 2014). Besides, there were increased expression of genes such as *MGST1* and *MRPS30-DT*, which were known to promote breast carcinoma cell growth and metastasis (Wu, et al., 2019; Zeng, et al., 2020). In IDC-inner, there were higher levels of tumor activity and lower levels of immune response. *LINC00052*, known to promote breast cancer cell proliferation by increasing signals of epidermal growth factor receptor (EGFR) such as *HER3* (Huang, et al., 2022; Salameh, et al., 2017; Xiong, et al., 2021), was overexpressed. Upregulation of *COX6C* and *FAM234B* implied higher levels of cellular respiration (Grzybowska-Szatkowska and Slaska, 2014) and lower immune response function (Lyu, et al., 2020), respectively.

Furthermore, we inferred developmental trajectories for the IDC and DCIS/LCIS regions. The starting points for calculating pseudo-time in IDC and DCIS/LCIS were the interior of the tumor regions (Fig. 6F). There were increased pseudo-time values as the region of the tumor moves outward, effectively mimicking the gradual progression of tumor development.

Moreover, to further evaluate the power of SECE in spatial domain identification, we tested 12 Human dorsolateral prefrontal cortex (DLPFC) datasets generated from the Visium platform. The original study (Maynard, et al., 2021) had manually annotated the spatial domains of these datasets, encompassing white matter and six cortical layers (Supplementary Figure S13, S14). Spatial domain identified by SECE exhibited the highest agreement with the original annotations. The median ARI for SECE was 0.58, surpassing the second and third ranked algorithms, STAGATE and SpatialPCA, which achieved median ARI values of 0.55 and 0.54, respectively. These findings highlighted the superior performance of SECE in accurately delineating the spatial domains within the low-resolution data.

## 4 Discussion

SECE captures both local and global relationships among spots and aggregates their information using expression similarity and spatial similarity, respectively. This approach enables precise spatial domain division and facilitates interpretable spatial embedding learning across diverse ST datasets. Moreover, The AE module that explicitly models gene expression counts enhances SECE’s ability to handle noisy data. Current methods use scaled values of highly variable genes (HVG) (e.g., GraphST, SpatialPCA, STAGATE, SpaceFlow) or employ dimensionality reduction techniques like principal component analysis (PCA) (e.g., BASS, SpaGCN, BayesSpace, conST) for expression features.

With the increasing captured area of ST data and advancements in resolution, there is a growing demand for computational methods to exhibit higher efficiency and scalability. We recorded the runtime and GPU memory consumption for each dataset (Supplementary Figure S15). For Slide-seqV2 hippocampus data and the Stereo-seq hemibrain data, which contained over 50,000 cells, SECE achieved a running time of less than 4.2 minutes while utilizing less than 5GB of GPU memory. These results demonstrated the superior computational efficiency and scalability of SECE when dealing with large-scale datasets.

While SECE has demonstrated notable performance, there are still several aspects that can be further enhanced. Firstly, we employ a pre-defined SSM to characterize global similarity, but exploring more flexible global correlation patterns could be advantageous. Secondly, we only utilized ST data as input, but incorporating matching histology data may provide additional benefits (Hu, et al., 2021). Although matching histology image data is currently only available on specific platforms like Visium, we can still leverage such images as optional supplementary information when available. Finally, integrating single-cell data with ST data can enhance the data quality of the latter, increase the throughput, or reduce the noise in the gene expression (Biancalani, et al., 2021). Therefore, incorporating single-cell data is another approach to improve spatial representation capabilities.

## Data availability

The datasets are available at Zenodo https://zenodo.org/record/8130682.

