## Supplementary Materials for "SECE: accurate identification of spatial domain by incorporating global spatial proximity and local expression proximity"

##### 1 Supplementary Notes

#### 1.1 ST datasets

***Mouse visual cortex***

Mouse visual cortex (Wang, et al., 2018) was generated from STARmap platform with 1,207 cells, 1,020 genes, and a sparsity of 76.88%. STARmap is a *in situ* sequencing-based ST method with single-molecule resolution. Despite its low gene throughput (160 to 1020 genes), it offers high sensitivity at single cell resolution with high efficiency and reproducibility. The dataset is available at <http://clarityresourcecenter.org/>.

***Mouse hippocampus dataset***

Mouse hippocampus dataset was generated from Slide-seqV2 (Stickels, et al., 2021) platform, with 53,208 cells and 23,264 genes from hippocampus, cortex, and thalamus, boasting a high sparsity of 98.19%. Slide-seqV2 offers transcriptome-wide sequencing with near-cellular resolution (10 μm). The dataset is available at <https://singlecell.broadinstitute.org/>.

***Mouse olfactory bulb***

Mouse olfactory bulb (Fu, et al., 2021) was generated from Stereo-seq (Chen, et al., 2022) platform, comprising 19,527 cells and 27,106 genes with 98.69% values were zero. Stereo-seq is an emerging technique for ST with genome-wide throughput and subcellular resolution. This method captures the expression profile and spatial coordination of each DNA nanoball (DNB) and employs image-based cell segmentation to segment single cells. It is available at <https://github.com/JinmiaoChenLab/SEDR_analyses>.

***Mouse hemibrain***

Mouse hemibrain was also generated from Stereo-seq (Chen, et al., 2022), which contained 50,140 cells and 25,879 genes with 96.94% values were 0. The dataset is available at <https://db.cngb.org/stomics/mosta/>.

***Human breast cancer***

Human breast cancer data was generated from Visium platform containing 3,798 spots and 24,923 genes, of which only 77.44% are zero values. Visium is the commercial version of Spatial transcriptomics (Stahl, et al., 2016) with a low resolution of 55 μm spots and 1-10 cells per spot (Larsson, et al., 2021). The dataset is available at <https://www.10xgenomics.com/resources/datasets>.

***Human dorsolateral prefrontal cortex***

Human dorsolateral prefrontal cortex (DLPFC) (Maynard, et al., 2021) dataset was also generated from Visium platform. 12 DLPFC slices were annotated manually and used as ground truth of spatial domain identification. The dataset is available within the spatialLIBD package (http:://spatial.libd.org/spatialLIBD).

***Preprocessing***

For Stereo-seq datasets, cells with an expression level below 200 were removed according to original studies (Chen, et al., 2022), then we filtered genes expressed in less than 20 cells. For other datasets, we screened spots with an expression level below 20 and genes that expressed less than 20 spots. The filtered expression matrix and its corresponding coordinates were input into SECE for analysis.

#### 1.2 Evaluation metrics

***Adjusted rand index (ARI)***

The ARI evaluates the degree of overlap between the two divisions which is formulated as:

$$ARI=\frac{\sum_{ij} \left( \begin{aligned} n_{ij} \\ 2 \end{aligned} \right)-[\sum_{i} \left( \begin{aligned} a_{i} \\ 2 \end{aligned} \right)\sum_{j} \left( \begin{aligned} b_{j} \\ 2 \end{aligned} \right)]/\left( \begin{aligned} N \\ 2 \end{aligned} \right)}{\frac{1}{2}[\sum_{i} \left( \begin{aligned} a_{i} \\ 2 \end{aligned} \right)+\sum_{j} \left( \begin{aligned} b_{j} \\ 2 \end{aligned} \right)]-[\sum_{i} \left( \begin{aligned} a_{i} \\ 2 \end{aligned} \right)\sum_{j} \left( \begin{aligned} b_{j} \\ 2 \end{aligned} \right)]/\left( \begin{aligned} N \\ 2 \end{aligned} \right)}$$

Where $N$ is the number of samples, $n_{ij},a_{i}$and $b_{j}$ are values from the contingency table. Specifically, $a_{i}$ represents the number of samples with the real category label $i$; $b_{j}$ represents the number of samples with the predicted label $j$; $n_{ij}$ represents the number of samples with the real category label $i$ and the predicted label $j$. In this paper, we utilize ARI to evaluate the consistency of the spatial domains identified by various methods with the ground truth domains. ARI ranges from -1 to 1, a greater value indicates better agreement with the true labels.

***Accuracy (ACC)***

The ACC evaluates the correctness of categories which is calculated as:

$$ACC=\frac{\sum_{i=1}^{N} \delta(r_{i}, o(s_{i}))}{N},$$

where $N$ is the number of samples, $r_{i}$ and $s_{i}$ are the true and predicted spatial domain label of spot $i$. $\delta$ is a function that can be defined as

$$\delta\left( x,y \right)=\left\{ \begin{aligned} 1, x=y \\ 0, x\neq y \end{aligned} \right.$$

$o$ is a mapping function that takes the real label $r_{i}$ as the reference label and then rearranges $s_{i}$ in the same arrangement, which is implemented using the classical Kuhn-Munkres algorithm (Munkres, 1957). ACC ranges from 0 to 1, a greater value indicates better performance.

***Average silhouette width (ASW)***

The ASW describes the degree of match between features and category labels. For every spot $i$，silhouette width $S\left( i \right)$is calculated as:

$$S\left( i \right)=\frac{b\left( i \right)-a(i)}{max\{a\left( i \right),b(i)\}},$$

where $a(i)$ is the average distance between $i$ and points in its cluster, $b(i)$ is the lowest average distance from $i$ to points in other clusters. In this paper, we use ASW to evaluate how well the SE obtained by various methods explain the known spatial layers. Distance is calculated by SE of various methods, and clusters are annotated spatial layers. ASW ranges from -1 to 1. A greater ASW indicates better SE learning.

***Local inverse Simpson’s index (******LISI)***

The $\mathrm{LISI}$ (Korsunsky, et al., 2019) measures the degree of local mixing to evaluate the level of spatial aggregation patterns. For each spot $i$，LISI can be formulated as:

$$LISI\left( i \right)=\frac{1}{\sum_{l\in L} p_{i}\left( l \right)},$$

where $p(l)$ is the probability that the spatial domain cluster label $l$ exists in the local neighborhood of spot $i$, and $L$ is the set of spatial domains. In this paper, we use LISI to evaluate the spatial aggregation degree of spatial domains. Local neighborhoods are generated by spatial location, cluster labels are those predicted by each algorithm. LISI value ranges from 0 to 1. A smaller LISI indicates better spatial aggregation patterns, i.e. less mixing of cluster labels within local spatial neighborhoods.

#### 1.3 Methods for comparison

We compared SECE with the existing spatial domain identification methods.

(1) BayesSpace implemented in the R package *BayesSpace* downloaded from <https://github.com/edward130603/BayesSpace>; (2) SpaGCN implemented in the Python package *SpaGCN* downloaded from <https://github.com/jianhuupenn/SpaGCN>*.* (3) STAGATE implemented in the Python package *STAGATE_pyG* downloaded from <https://github.com/QIFEIDKN/STAGATE>. (4) BASS implemented in the R package *BASS* from <https://github.com/zhengli09/BASS>. (5) SpaceFlow implemented in the Python package *SpaceFlow* from <https://github.com/hongleir/SpaceFlow>. (6) GraphST implemented in the Python package *GraphST* from <https://github.com/JinmiaoChenLab/GraphST>. (7) SpatialPCA implemented in the R package *SpatialPCA* from <https://github.com/shangll123/SpatialPCA>. (8) conST. We referred to <https://github.com/ys-zong/conST> to ran conST.

Chen, A.*, et al.* Spatiotemporal transcriptomic atlas of mouse organogenesis using DNA nanoball-patterned arrays. *Cell* 2022;185(10):1777-1792 e1721.

Fu, H.*, et al.* Unsupervised Spatially Embedded Deep Representation of Spatial Transcriptomics. *bioRxiv* 2021:2021.2006.2015.448542.

Korsunsky, I.*, et al.* Fast, sensitive and accurate integration of single-cell data with Harmony. *Nat Methods* 2019;16(12):1289-1296.

Larsson, L., Frisen, J. and Lundeberg, J. Spatially resolved transcriptomics adds a new dimension to genomics. *Nat Methods* 2021;18(1):15-18.

Maynard, K.R.*, et al.* Transcriptome-scale spatial gene expression in the human dorsolateral prefrontal cortex. *Nat Neurosci* 2021;24(3):425-436.

Munkres, J. Algorithms for the Assignment and Transportation Problems. *Journal of the Society for Industrial and Applied Mathematics* 1957;5(1):32-38.

Stahl, P.L.*, et al.* Visualization and analysis of gene expression in tissue sections by spatial transcriptomics. *Science* 2016;353(6294):78-82.

Stickels, R.R.*, et al.* Highly sensitive spatial transcriptomics at near-cellular resolution with Slide-seqV2. *Nat Biotechnol* 2021;39(3):313-319.

Wang, X.*, et al.* Three-dimensional intact-tissue sequencing of single-cell transcriptional states. *Science* 2018;361(6400).

### 2 Supplementary Figures

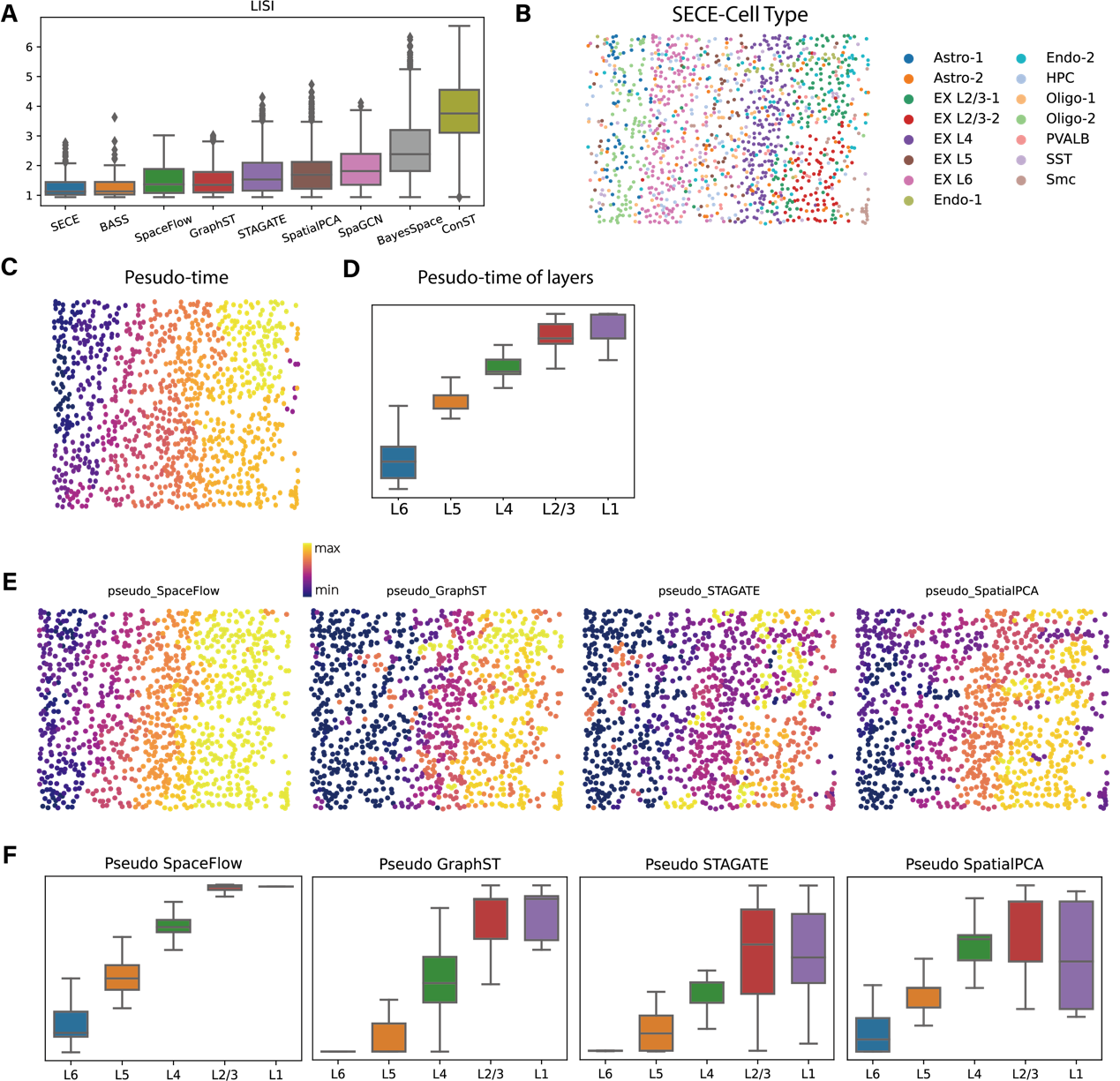

**Fig. S1.** **Trajectory inference on mouse visual cortex STARmap data, related to Fig.2. (A)** Boxplot of LISI measuring spatial aggregation of domains identified by different methods. **(B)** Cell type annotation. Astro, astrocytes; Oligo, oligodendrocytes; EX, excitatory neurons; IN Neuron, inhibitory neurons; Endo, endothelial cells; SMC, smooth muscle cells. **(C)** Pseudo-time of each cell calculated by Monocle3 based on SECE embeddings. **(D)** Pseudo-time of cells in each cortical layer based on SECE. (**E)** Pseudo-time of each cell calculated by Monocle3 based on SpaceFlow, GraphST, STAGATE, and SpatialPCA embeddings. (**F)** Pseudo-time of cells in each cortical layer based on SpaceFlow, GraphST, STAGATE and SpatialPCA.

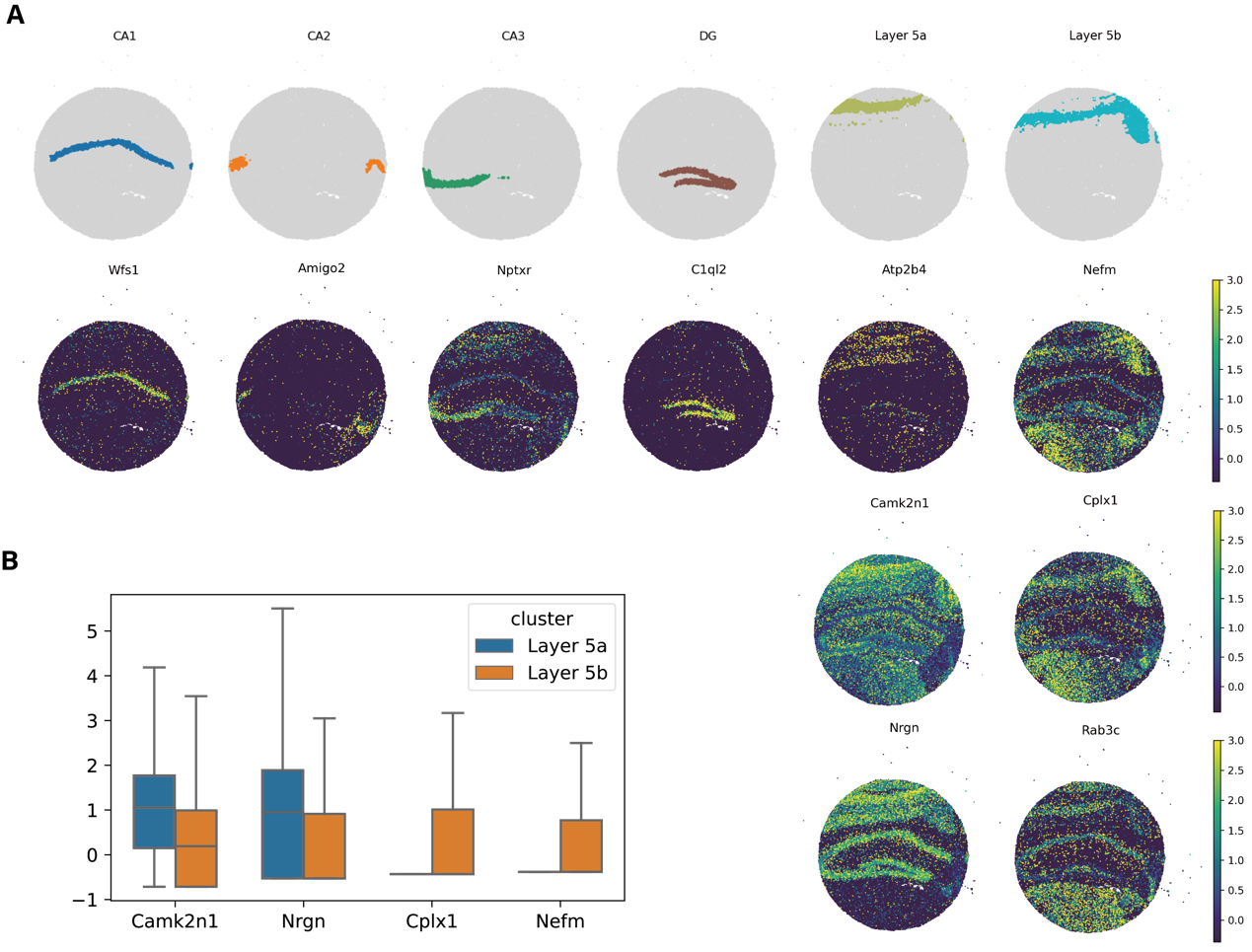

**Fig. S2.** **Marker of spatial domains identified by SECE on mouse hippocampus Slide-seqV2 data, related to Fig.3. (A)** Spatial visualization of CA1, CA2, and CA3 domains identified by SECE (Top) and the corresponding marker genes (Bottom). **(B)** Boxplots showing expression levels of *Camk2n1*, *Nrgn*, *Cplx1*, *Nefm* in Layer 5a and Layer 5b.

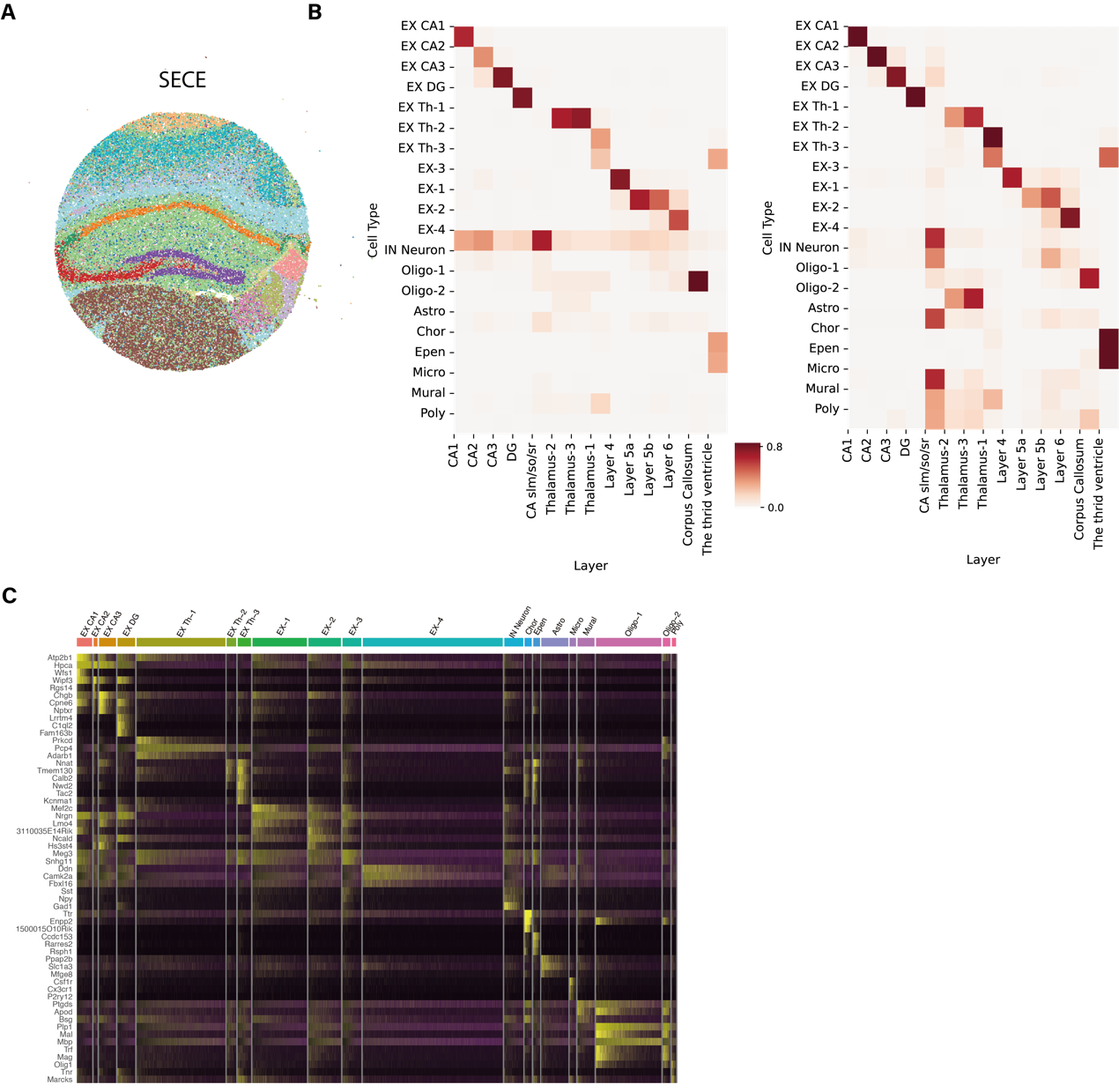

**Fig. S3. Cell type composition of domains in mouse hippocampus Slide-seqV2 data, related to Fig.3. (A)** Spatial visualization of cell types. **(B)** Left: Heatmap showing proportion of each cell type across spatial domains identified by SECE. The color represents the proportion of each cell type contained in each spatial region. The sum of the proportions of each spatial domain is 1. Right: Heatmap showing distribution of each cell type across spatial domains. The color represents what proportion of each cell type is distributed in that spatial region. The sum of the proportions of each cell type is 1. **(C)** Gene expression heatmaps for cell type clusters. Each gene was centered and standardized across all the cells. For each cell type cluster, gene expression of the top three differentially expressed (DE) genes are displayed, where DE genes were identified using the Wilcoxon rank-sum test contrasting each cluster of cells against all the remaining cells. Cell type clusters were annotated with specific cell types by comparing the identified DE genes with previously known cell type marker genes.

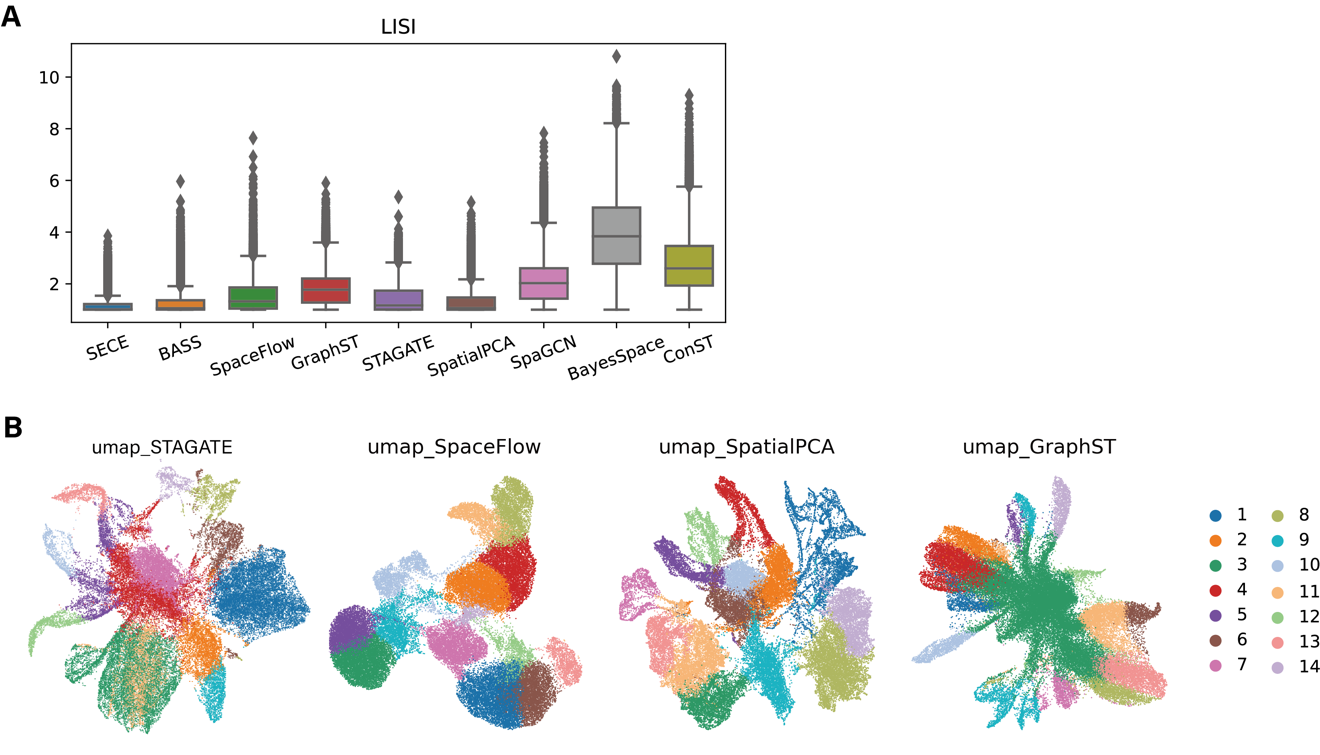

**Fig. S4. Spatial domains of mouse hippocampus Slide-seqV2 data, related to Fig.3. (A)** Boxplot of LISI measuring spatial aggregation of domains identified by different methods. **(B)** UMAP visualizations generated by SpaceFlow, GraphST, STAGATE, and SpatialPCA, colored by corresponding domains.

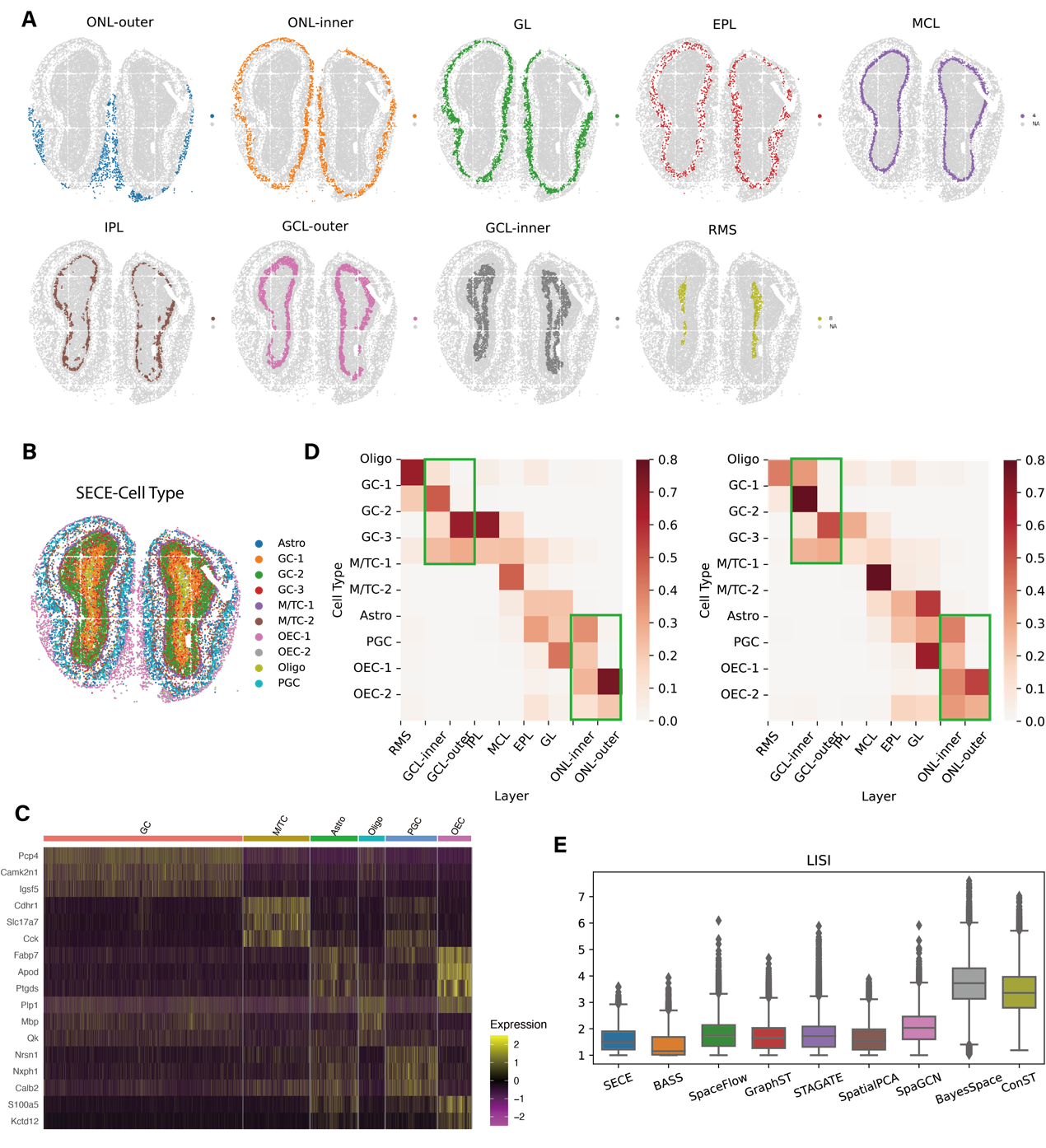

**Fig. S5. Spatial domain identification of olfactory bulb, related to Fig.4. (A)** Separate spatial visualization for each identified layer. **(B)** Spatial visualization of cell types. GC, Granule cells; M/TC, Mitral and tufted cells; PGC, Periglomerular cells; OEC, Olfactory ensheathing cells. **(C)** Gene expression heatmaps for cell type clusters. Each gene was centered and standardized across all the cells. For each cell type cluster, gene expression of the top three differentially expressed (DE) genes are displayed, where DE genes were identified using the Wilcoxon rank-sum test contrasting each cluster of cells against all the remaining cells. Cell type clusters were annotated with specific cell types by comparing the identified DE genes with previously known cell type marker genes. **(D)** Left: Heatmap showing proportion of each cell type across spatial domains. The color represents the proportion of each cell type contained in each spatial region. The sum of the proportions of each spatial domain is 1. Right: Heatmap showing distribution of cell types across spatial domains. The color represents what proportion of each cell type is distributed in that spatial region. The sum of the proportions of each cell type is 1. **(E)** Boxplot of LISI measuring spatial aggregation of domains identified by different methods.

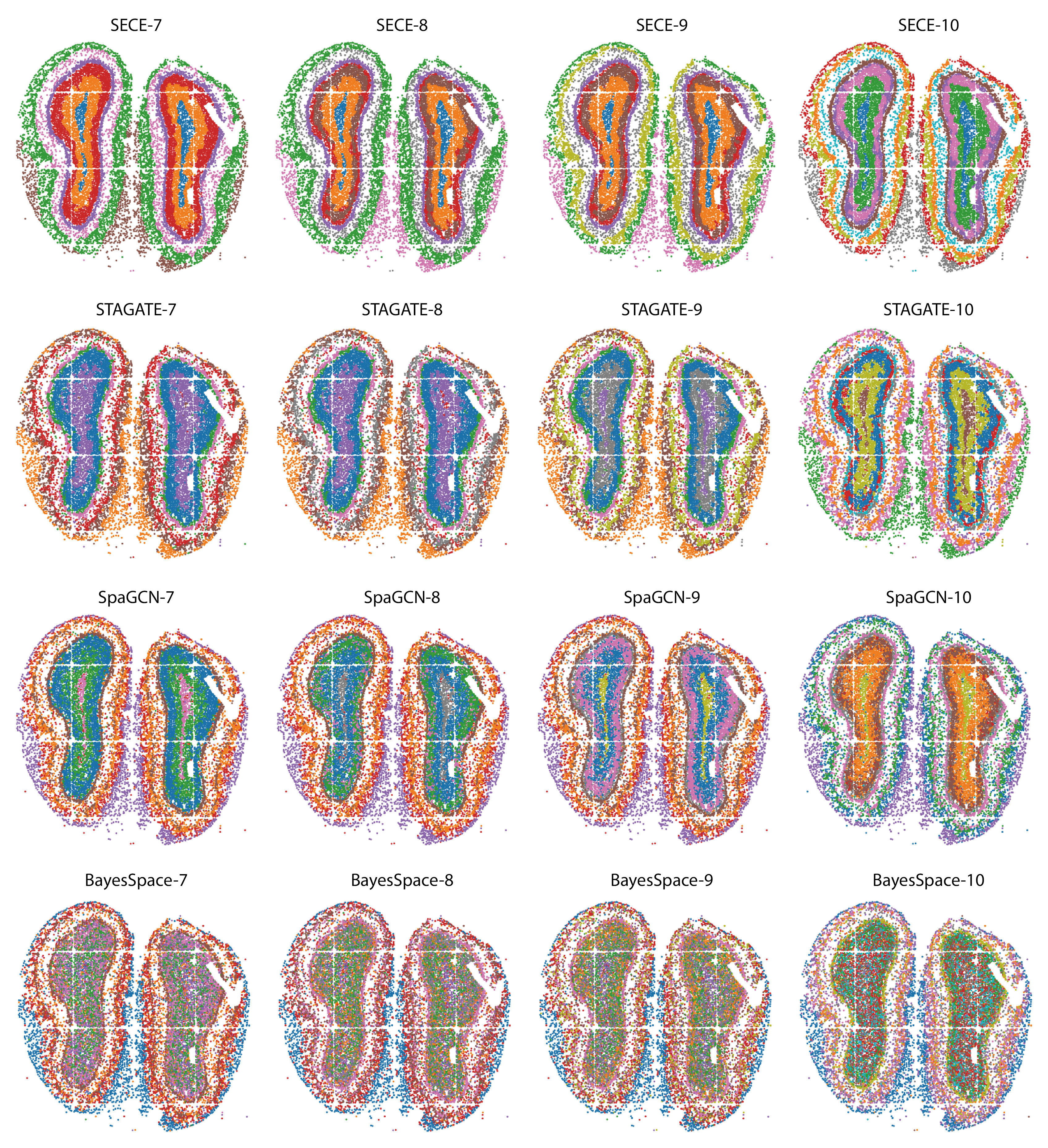

**Fig. S6. Different number (7, 8, 9, 10) of spatial domains identified by the SECE, STAGATE, SpaGCN and BayesSpace in the Stereo-seq mouse olfactory bulb data.**

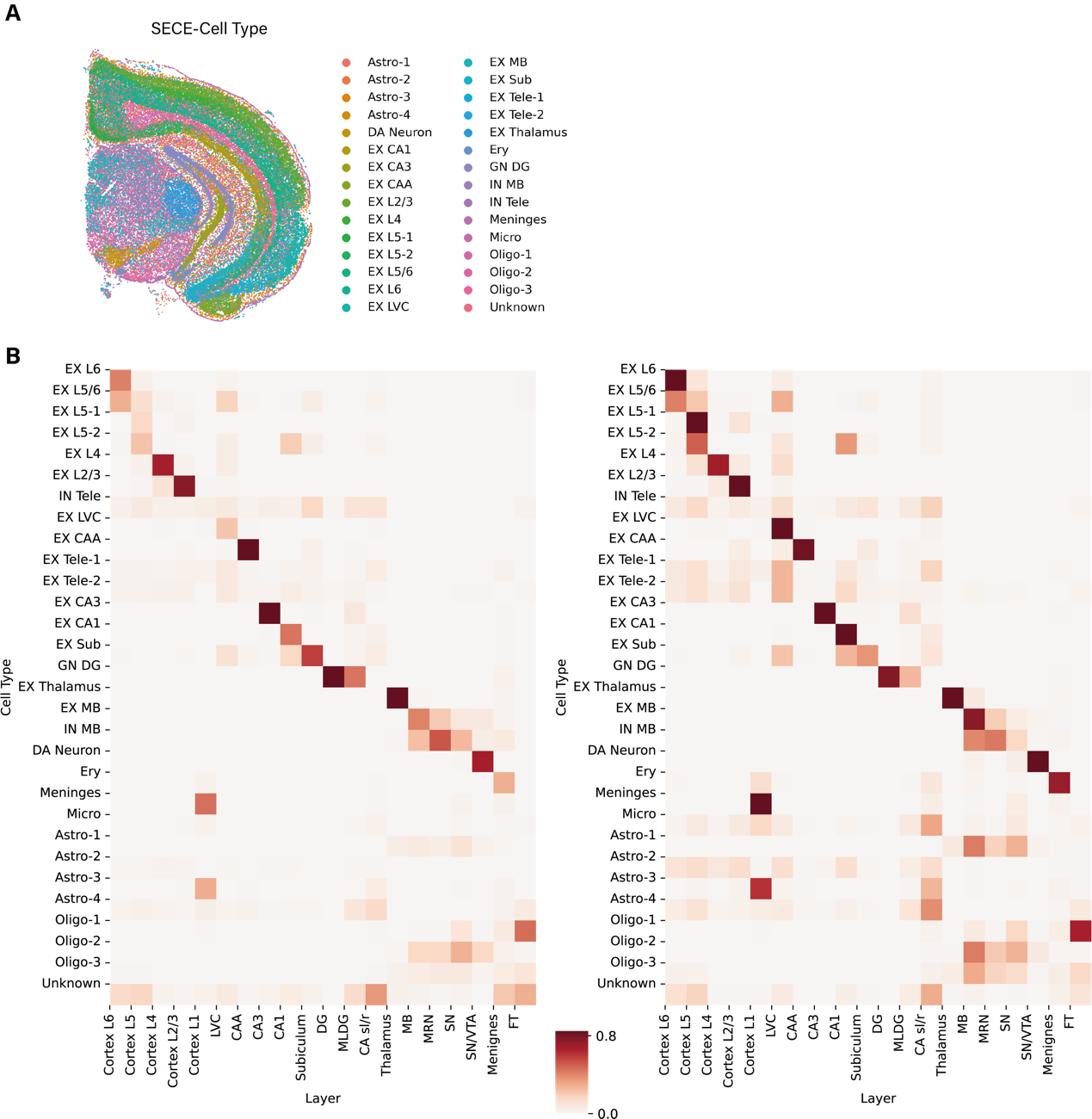

**Fig. S7. Relationship between cell types and spatial regions identified by SECE of mouse brain Stereo-seq data, related to Fig.5.** **(A)** Spatial visualization of cell types. **(B)** Left：Heatmap showing proportion of each cell type across spatial domains identified by SECE. The color represents the proportion of each cell type contained in each spatial region. The sum of the proportions of each spatial domain is 1. Right: Heatmap showing distribution of each cell type across spatial domains identified by SECE. The color represents what proportion of each cell type is distributed in that spatial region. The sum of the proportions of each cell type is 1.

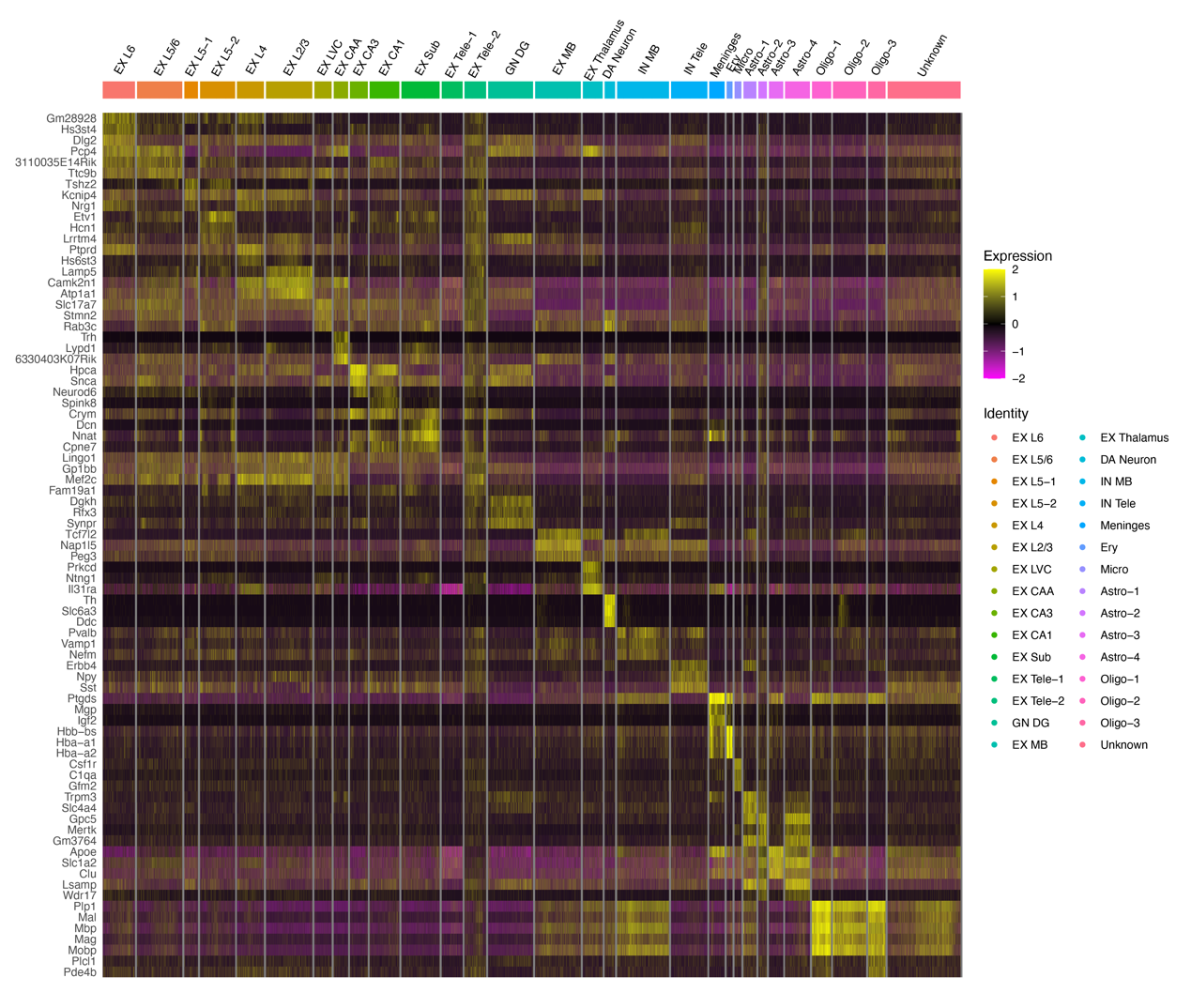

**Fig. S8. Gene expression heatmaps of cell type clusters in mouse brain Stereo-seq data, related to Fig.5.** Each gene was centered and standardized across all the cells. For each cell type cluster, gene expression of the top three differentially expressed (DE) genes are displayed, where DE genes were identified using the Wilcoxon rank-sum test contrasting each cluster of cells against all the remaining cells. Cell type clusters were annotated with specific cell types by comparing the identified DE genes with previously known cell type marker genes.

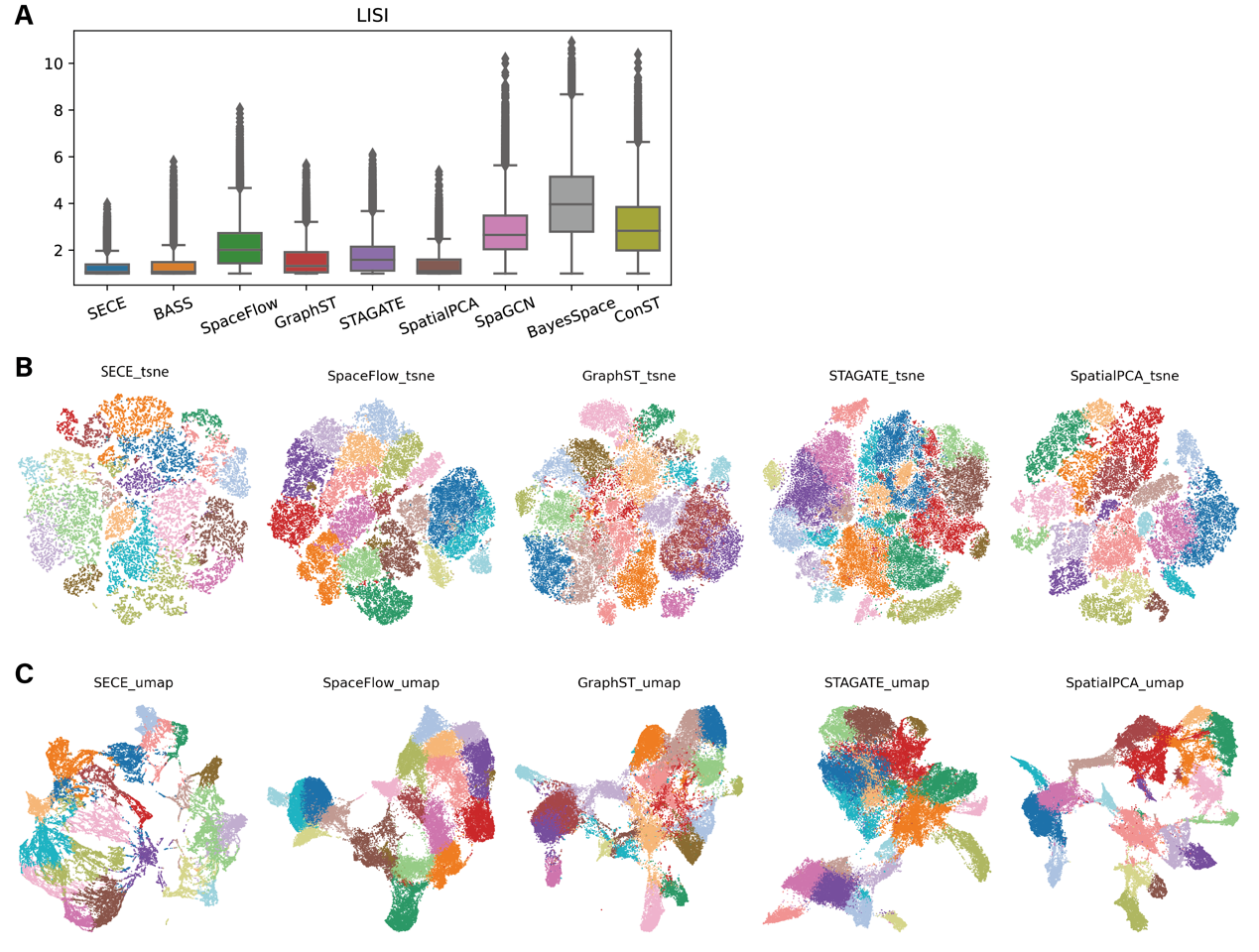

**Fig. S9. Spatial domain identification of mouse brain Stereo-seq data, related to Fig.5.** (**A)** Boxplot of LISI measuring spatial aggregation of domains identified by different methods. (**B)** t-SNE visualizations generated by SECE, SpaceFlow, GraphST, STAGATE, and SpatialPCA embeddings, colored by corresponding spatial domains. **(C)** UMAP visualizations generated by SECE, SpaceFlow, GraphST, STAGATE, and SpatialPCA embeddings, colored by corresponding spatial domains.

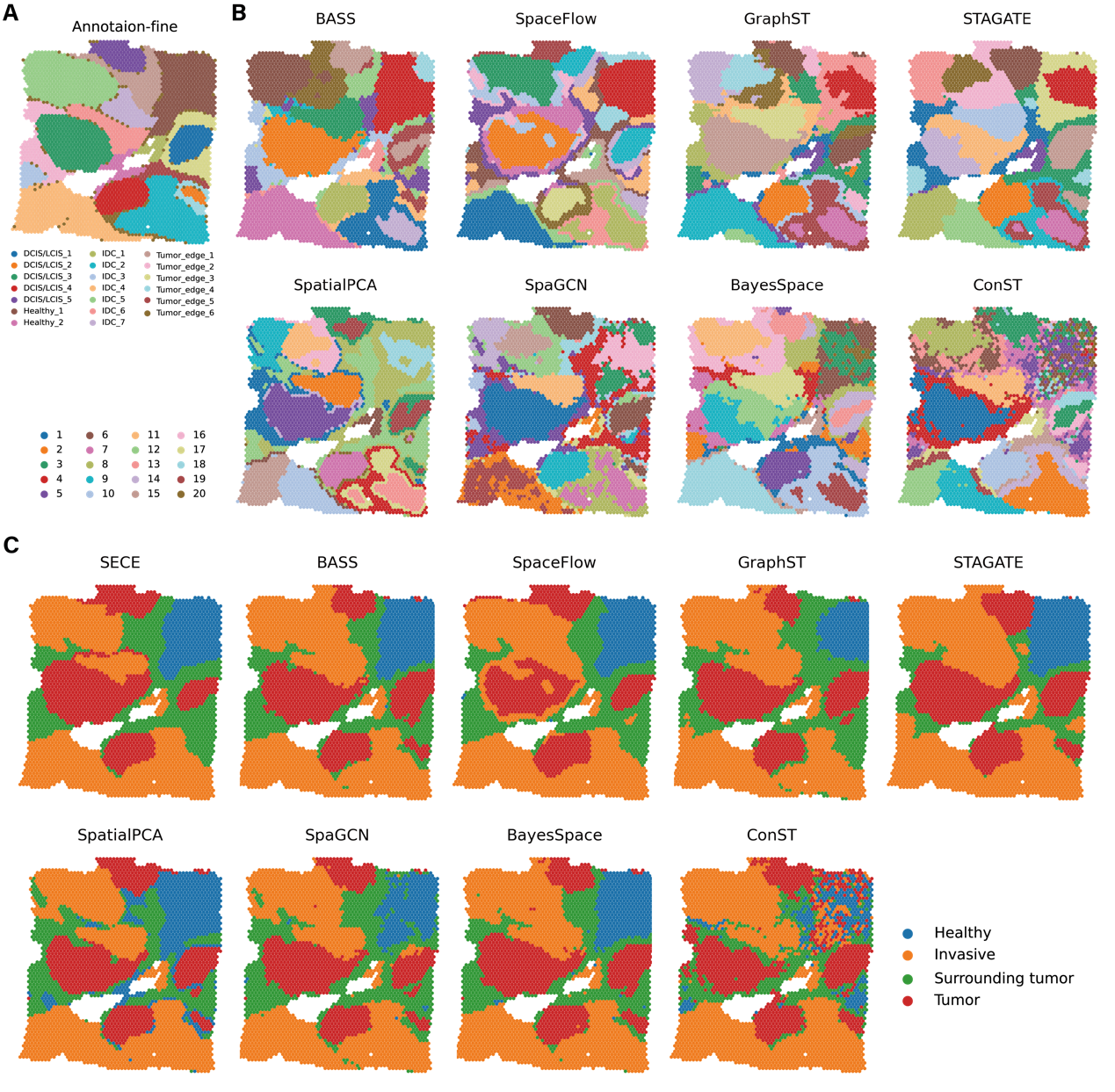

**Fig. S10. Spatial domain identification of breast cancer data, related to Fig.6. (A)** Pathology annotation of the tissue section from the original study. **(B)** Spatial regions identified by different methods. **(C)** Annotation of Spatial regions identified by different methods.

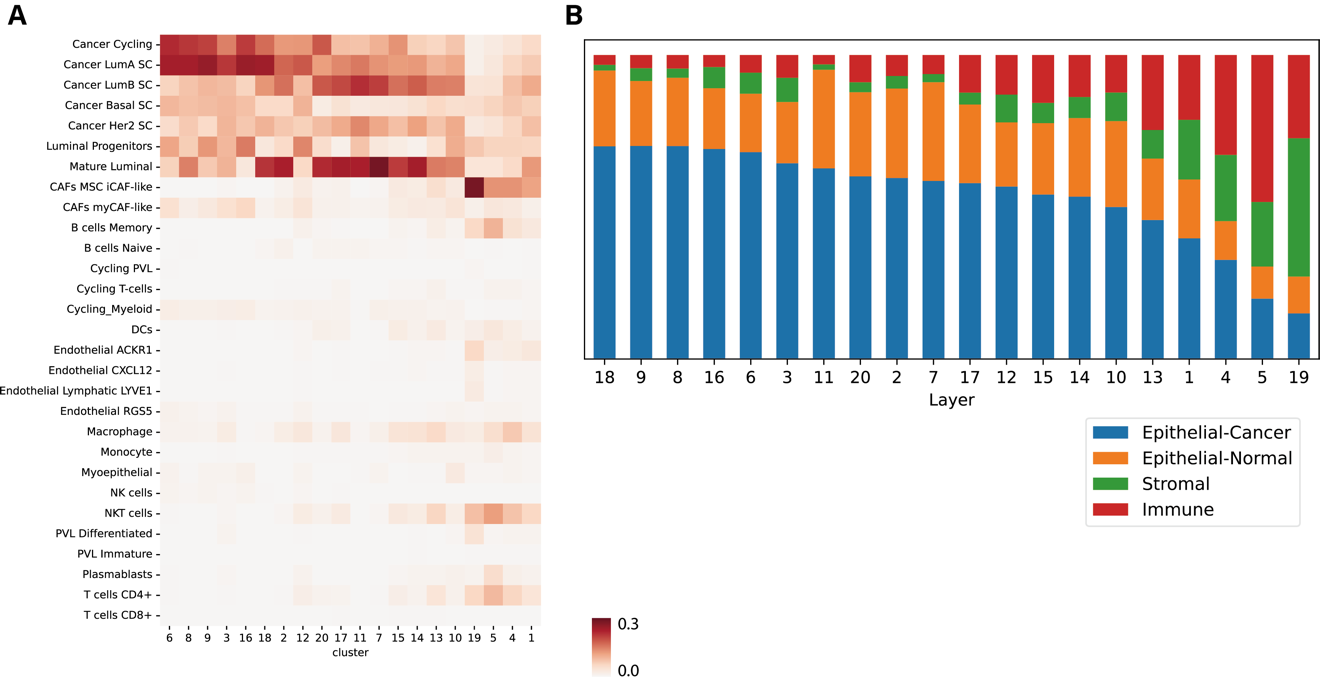

**Fig. S11. Cell type composition in spatial regions identified by SECE, related to Fig.6. (A)** Heatmap showing proportion of each cell type across spatial domains identified by SECE, with cell types inferred by cell2location. (**B)** Stacked bar chart showing proportion of summarized cell types across spatial domains. The four cell types are consolidated from the cell2location predictions.

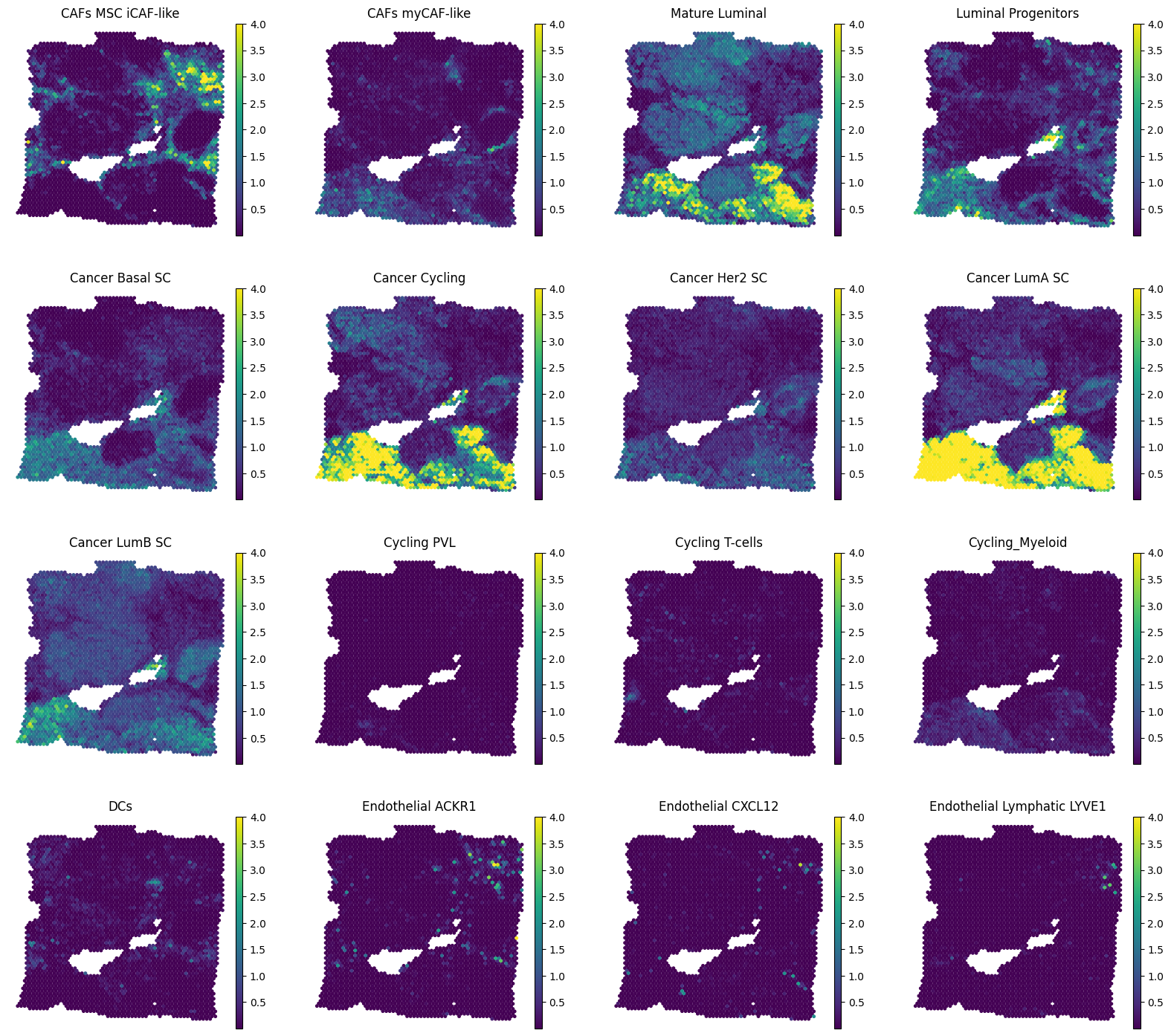

See next page

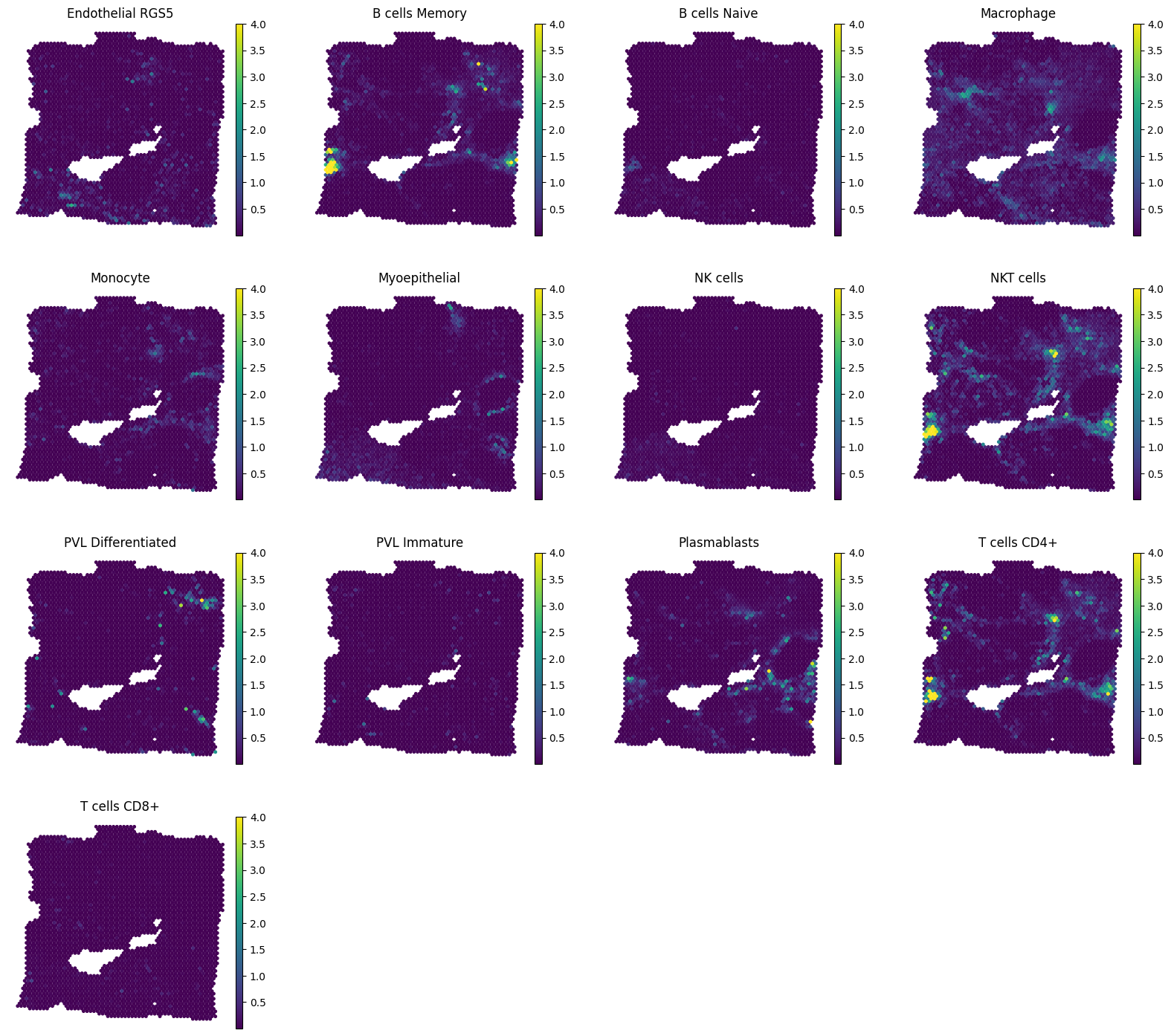

**Fig. S12. Number of cells per spot for each cell type inferred by cell2location in human breast cancer Visium data.**

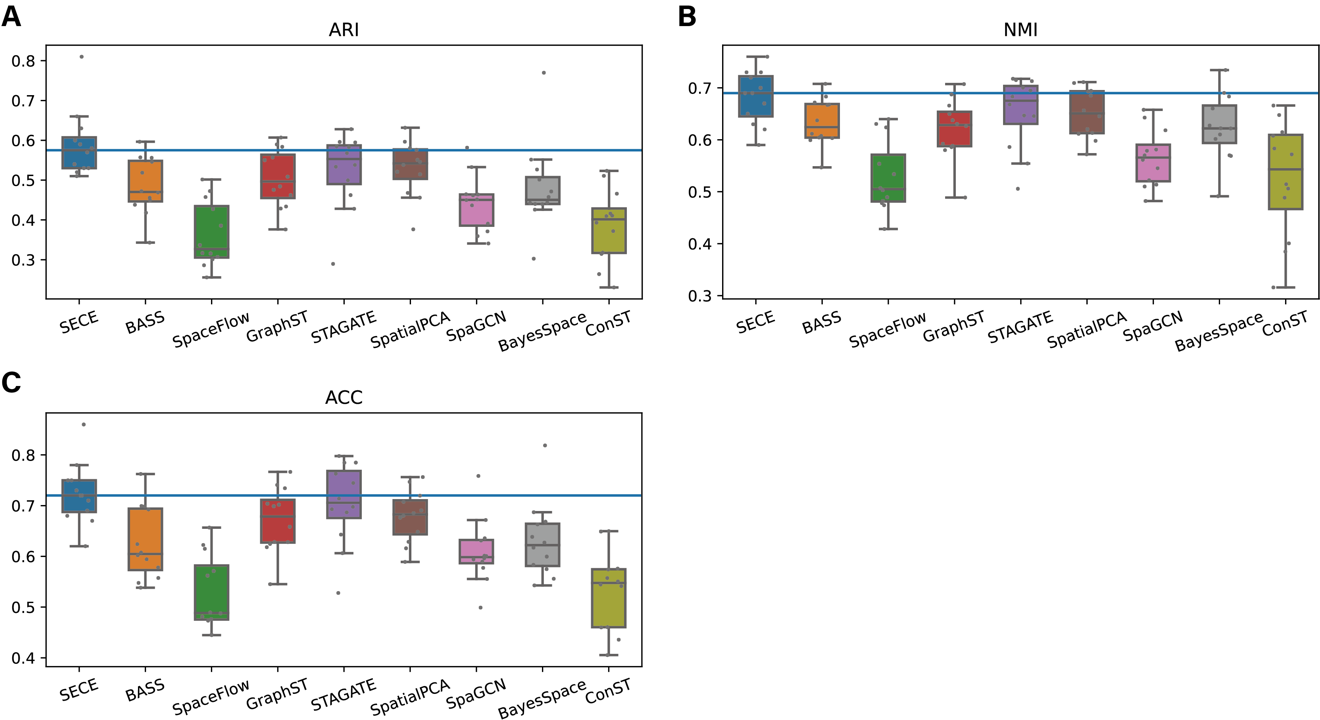

**Fig. S13. Comparison of spatial domain identification on 12 sections of the DLPFC dataset. (A)** Boxplot of clustering accuracy across sections in terms of ARI. **(B)** Boxplot of clustering accuracy across sections in terms of NMI. **(C)** Boxplot of clustering accuracy across sections in terms of ACC.

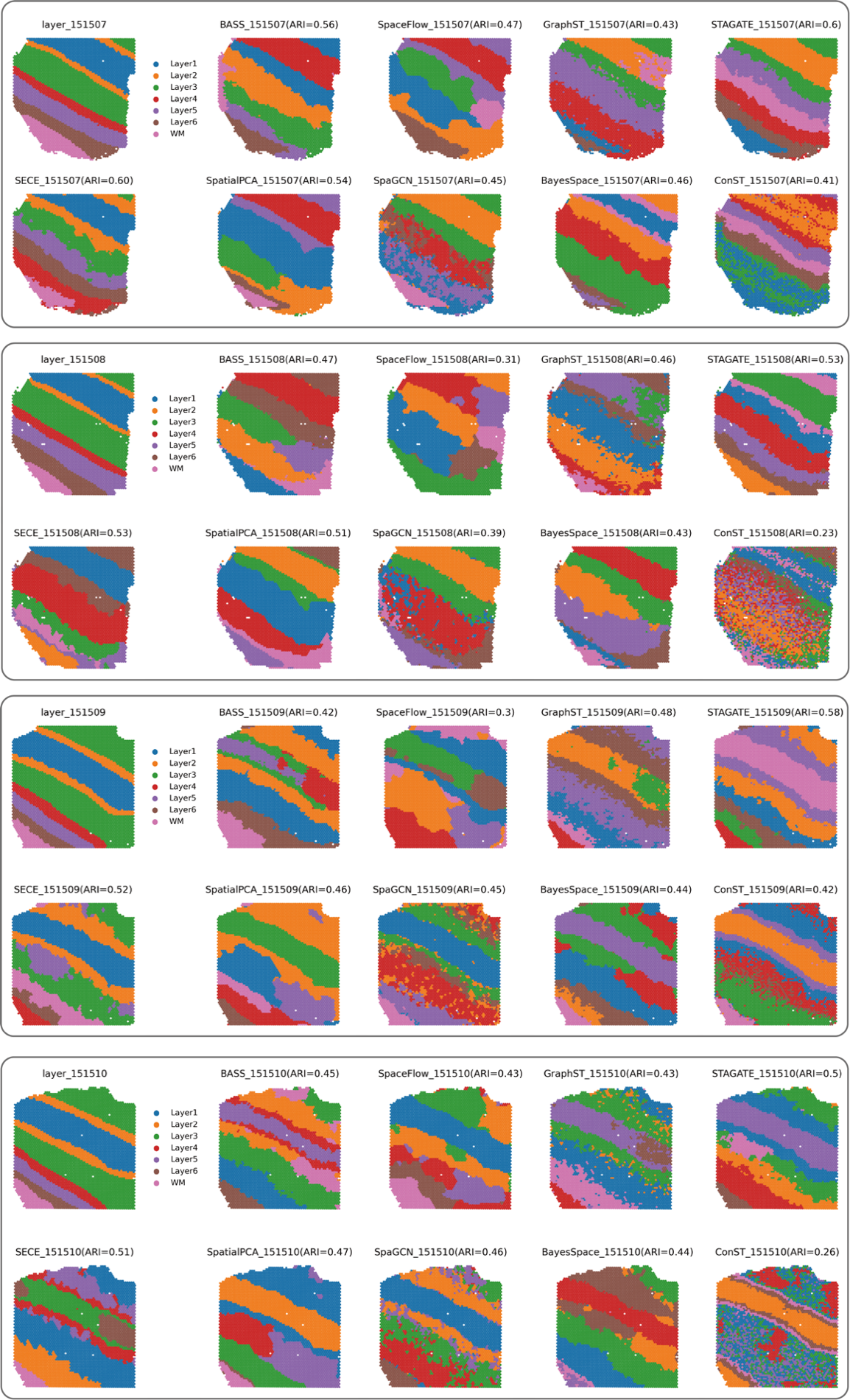

See next page

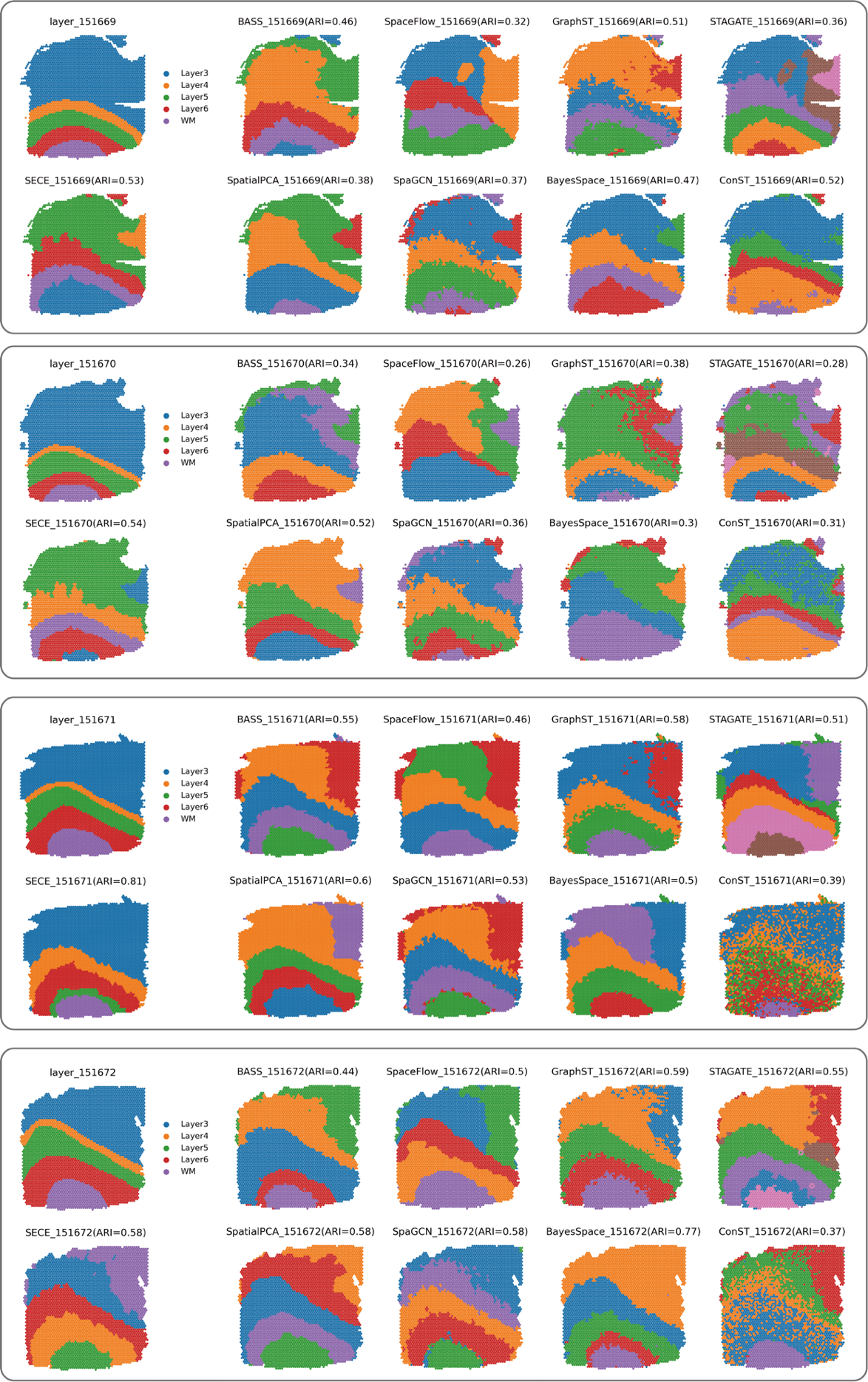

See next page

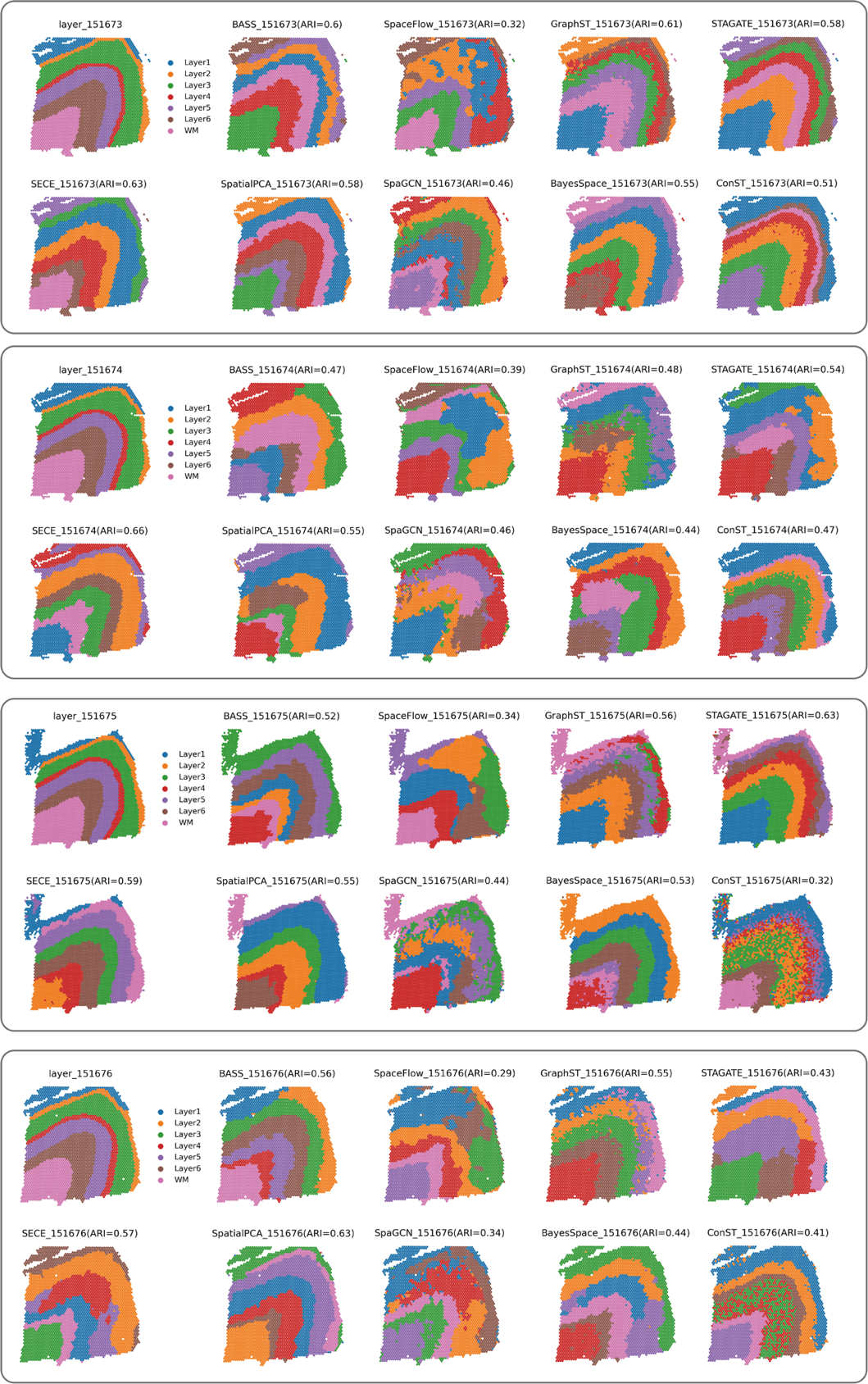

**Fig. S14. Spatial domains identified by SECE, BASS, SpaceFlow, GraphST, STAGATE, SpatialPCA, SpaGCN, BayesSpace, and conST, and manual annotation in 12 sections of the DLPFC dataset.**

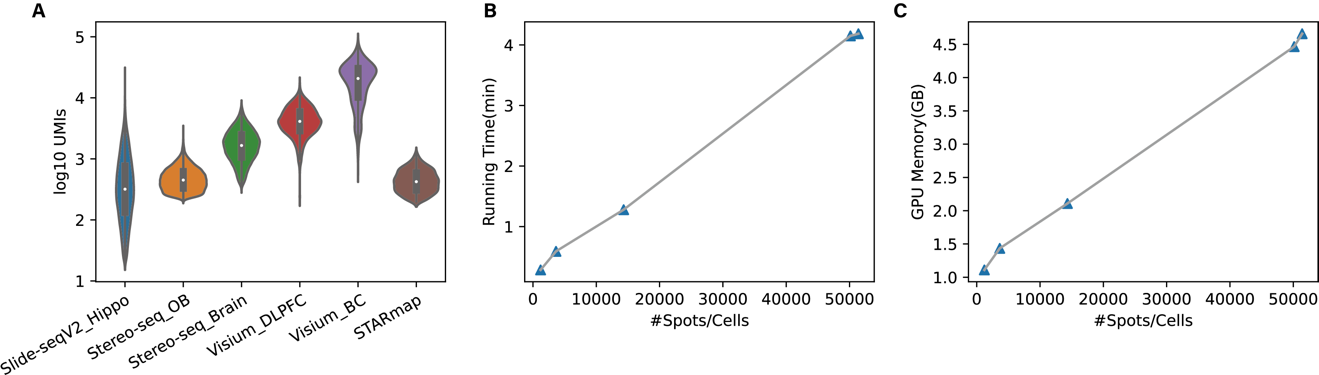

**Fig. S15. Datasets used by SECE and their running information. (A)** Number of total UMIs per spot in each dataset. Running time **(B)** and GPU memory usage **(C)** on dataset with different numbers of spots based on NVIDIA® Tesla® V100 GPU.
